# Long-read Assays Shed New Light on the Transcriptome Complexity of a Viral Pathogen and on Virus-Host Interaction

**DOI:** 10.1101/2020.01.27.921056

**Authors:** Dóra Tombácz, István Prazsák, Zoltán Maróti, Norbert Moldován, Zsolt Csabai, Zsolt Balázs, Béla Dénes, Tibor Kalmár, Michael Snyder, Zsolt Boldogkői

**Affiliations:** Department of Medical Biology, Faculty of Medicine, University of Szeged, Szeged, Hungary; Department of Pediatrics, Faculty of Medicine, University of Szeged, Korányi fasor 14-15., Szeged, H-6720, Hungary; Veterinary Diagnostic Directorate of the National Food Chain Safety Office, Budapest, Hungary; Department of Genetics, School of Medicine, Stanford University, Stanford, California, USA

**Keywords:** poxvirus, vaccinia virus, transcriptome, long-read sequencing, Pacific Biosciences, RSII sequencing, Sequel system, Oxford Nanopore Technologies, MinION system, direct RNA sequencing

## Abstract

Characterization of global transcriptomes using conventional short-read sequencing is challenging because of the insensitivity of these platforms to transcripts isoforms, multigenic RNA molecules, and transcriptional overlaps, etc. Long-read sequencing (LRS) can overcome these limitations by reading full-length transcripts. Employment of these technologies has led to the redefinition of transcriptional complexities in reported organisms. In this study, we applied LRS platforms from Pacific Biosciences and Oxford Nanopore Technologies to profile the dynamic vaccinia virus (VACV) transcriptome and assess the effect of viral infection on host gene expression. We performed cDNA and direct RNA sequencing analyses and revealed an extremely complex transcriptional landscape of this virus. In particular, VACV genes produce large numbers of transcript isoforms that vary in their start and termination sites. A significant fraction of VACV transcripts start or end within coding regions of neighboring genes. We distinguished five classes of host genes according to their temporal responses to viral infection. This study provides novel insights into the transcriptomic profile of a viral pathogen and the effect of the virus on host gene expression.

**Author Summary:** Viral transcriptomes that are determined using conventional (first- and second-generation) sequencing techniques are incomplete because these platforms are inefficient or fail to distinguish between types of transcripts and transcript isoforms. In particular, conventional sequencing techniques fail to distinguish between parallel overlapping transcripts, including alternative polycistronic transcripts, transcriptional start site (TSS) and transcriptional end site (TES) isoforms, and splice variants and RNA molecules that are produced by transcriptional read-throughs. Long-read sequencing (LRS) can provide complete sets of RNA molecules, and can therefore be used to assemble complete transcriptome atlases of organisms. Although vaccinia virus (VACV) does not produce spliced RNAs, its transcriptome contains large numbers of TSSs and TESs for individual viral genes and has a high degree of polycistronism, together leading to enormous complexity. In this study, we applied single molecule real-time and nanopore-based cDNA and direct-RNA sequencing methods to investigate transcripts of VACV and the host organism.

## Introduction

Members of the *Poxviridae* family infect various vertebrate and invertebrate host species^1^. Among these, variola virus is the causative agent of smallpox^2^, but was extirpated in 1977 by a collaborative vaccination program using the closely related vaccinia (cowpox) virus (VACV) as a live vaccine^3^. VACV has roughly 90% sequence homology with the variola and is the prototype member of the genus of orthopoxviruses. VACV virion contains 195 kilobase pairs (kbp) of double-stranded DNA comprising at least 200 open reading frames (ORFs). Twelve of which are present in terminal repeats^4–6^. This viral genome encodes enzymes for DNA and RNA syntheses and transcription factors, and for enzymes that cap^7^ and polyadenylate^8^ RNA molecules, thus allowing VACV to replicate in the cytoplasm. Viral gene expression is regulated by stage-specific transcription factors that differentially recognize the promoters of early (E), intermediate (I), and late (L) genes^9–12^. The complete transcription machinery is packaged in the VACV virion, and therefore E genes can be expressed immediately after entering the cell, when the viral genome is still encapsidated^5^. Subsequently, DNA replication is followed by the synthesis of mRNAs from I- and then L-classes of genes. Synthesis of I mRNAs is dependent on *de novo* expression of E viral proteins, whereas synthesis of mRNAs from L genes requires the expression of certain E and I genes. During the last stage of infection, newly assembled virus particles egress from host cells. E genes encode proteins that synthesize DNA and RNA molecules, and others that play roles in virus-host interactions, whereas post-replicative (PR) genes, including I and L genes, mainly specify structural elements of the virus^13^. Kinetic classification of VACV genes was previously based on the strong preference for co-localization^14^: Specifically, E genes are reportedly clustered near genomic termini and are transcribed in the same direction^15^, whereas the I and L genes are clustered at the central part of the viral DNA and predominantly have the same orientations^16^. Using genome tiling arrays, Assarsson and colleagues demonstrated that 35 viral genes are expressed during the immediate-early (IE) stage of VACV replication^14^. Although this terminology has not become widely accepted, in other DNA viruses, such as herpesviruses^17^ and baculoviruses^18^, IE RNAs are expressed in the absence of *de novo* protein synthesis, whereas E and L RNAs are dependent on the synthesis of IE proteins. Thus, according to this classification, all VACV E transcripts should belong to the IE kinetic class^19^. Alternatively, however, Yang and colleagues categorized two early classes (E1.1 and E1.2) of VACV mRNAs^20^, and defined the ensuing expression kinetics according to higher affinity of transcription factor binding sites for E1.1 than for E1.2 promoters. Accordingly, the three classes of genes have distinctive promoters^11^, and the consensus transcription termination sequence of E transcripts (UUUUUNU) is non-functional in PR transcripts, which mostly have polymorphic 3’ ends^21–22^. Many PR and a few E transcripts have 5’ poly(A) tails (PAT) downstream of the cap structure, which was supposed to be generated by slippage of RNA polymerase on adjacent T’s^23^. In addition to genome tiling analyses^24^, VACV transcriptome analyses have been performed using RNA-Seq^20^, ribosome profiling^13, 25^, and microarrays^26–27^. Yang and colleagues mapped 118 E genes and 93 PR genes using a short-read sequencing (SRS) technique^15^, and in a later study, they distinguished 53 I and 38 L genes among the PR genes^16^. Ribosome profiling analysis also confirmed canonical translational initiation sites (TISs) and demonstrated that additional TISs occur mostly within the ORFs. These TISs were also detected in 5’-untranslated regions (UTRs), and intergenic regions, and even on the complementary DNA strands^13^. However, because most of these ORFs were short, the authors doubted their biological relevance. In a study by Yang and colleagues, transcript abundance was highly correlated with ribosome-protected reads, suggesting that translation is largely regulated by mRNA abundance^13^. The techniques used in the studies cited above cannot detect full-length RNA molecules, and hence fail to provide a comprehensive understanding of the viral transcriptome. Although deep SRS provides satisfactory sequencing depth and coverage for global transcriptome profiling, the resulting RNA molecule assemblies are incomplete, resulting unreliable annotations. The major limitations in VACV transcriptome profiling relate to multiple transcriptional read-throughs that generate a meshwork of overlapping RNAs and to the large variation in the transcript termini, including the transcription start sites (TSSs) and the transcriptional end sites (TESs), which can even be located within the coding sequences of the genes. Hence, conventional techniques that fail to read entire RNA molecules considerably underestimated the complexity of the poxvirus transcriptome. We used the Pacific Biosciences (PacBio) and Oxford Nanopore Technologies (ONT) long-read sequencing (LRS) platforms to profile the transcriptomes of herpesviruses, including pseudorabies virus^28–29^, herpes simplex virus type 1^30–31^, varicella-zoster virus^32^, and human cytomegalovirus^33–34^. These analyses sensitively identified polycistronic RNA molecules, transcript isoforms, and transcriptional overlaps, and facilitated the kinetic characterization of the viral transcripts^35–36^.

Herein, we report analyses from PacBio RSII and Sequel platforms, and from ONT MinION platform for cDNA, direct RNA (dRNA), and Cap-sequencing. Together, these techniques revealed the dynamic transcriptome of the Western Reserve (WR) strain of VACV. We achieved high coverage of sequencing reads and determined TSSs and TESs of RNA molecules at single-base resolution. We introduce the concepts of regular and chaotic genomic regions, and thereby indicate that a large variety of transcripts are produced from each ORF with a high heterogeneous TSSs and TESs at certain genomic positions, especially those containing I and L genes. The large diversity of initiation and termination of transcription creates considerable difficulty in identifying individual RNA molecules. Finally, we also evaluated temporal impacts of VACV infection on the host cell transcriptome.

## Results

### Long-read transcriptome sequencing of vaccinia virus and host cells

To investigate the transcriptomes of VACV and CV-1 host cells, we performed polyadenylation sequencing techniques using PacBio isoform sequencing (Iso-Seq) template preparation protocol for RSII and Sequel platforms, and using the ONT MinION device to sequence cDNAs and RNAs. For ONT sequencing, we used the company’s own library preparation approach (1D-Seq), or the Teloprime Cap-selection protocol (Cap-Seq) from Lexogen, which was adapted for the MinION sequencer. The obtained datasets were then used to define and validate viral TSS and TES coordinates using the LoRTIA pipeline (https://github.com/zsolt-balazs/LoRTIA) that was developed by our research group^37^. The workflow for library preparation and analysis is shown in **Fig 1**. Apart from dRNA sequencing, the techniques were based on polymerase chain reaction (PCR)-based amplification prior to sequencing. The dRNA-Seq tool offers an alternative to cDNA sequencing because it lacks the recoding and amplification biases inherent in reverse transcription (RT) and PCR-based methods. We also used random primer-based RT for some ONT sequencing analyses. All LRS methods achieved high data coverage and determined both TSSs and TESs of viral transcripts with base-pair precision. LRS can be used to detect individual RNA molecules and identify alternatively transcribed and processed RNA molecules, polycistronic transcripts, and transcriptional overlaps. Despite this straightforward utility, identification of transcript ends (especially, TSSs) was extremely challenging due to the tremendous complexity of the VACV transcriptome. The amplified SMRT Iso-Seq technique utilizes a switching mechanism at the 5’ end of the RNA template, thereby generating full-length cDNAs^38^. An advantageous feature of the PacBio technique is that any error that arises can be easily corrected due to its high consensus accuracy^39^. In this study, we used a workflow and pipeline for transcriptome profiling of long-read RNA sequencing data that was developed in our laboratory^31, 33, 40^. In total, about 1,115,000 reads of inserts (ROIs) were generated using the PacBio platform (**S1 Table**). In addition to PATs, ROIs can potentially be produced nonspecifically from A-rich regions of the RNA molecules^37^. Thus, we excluded these products from further analysis using the LoRTIA toolkit. Non-specific binding of the adapter and PCR primer sequences to cDNAs can also lead to false positives and we excluded these artifacts using bioinformatic filtering. We also used the LoRTIA toolkit to confirm that the 5’ and 3’ termini of the sequencing reads represented real TSSs and TESs of viral transcripts. When two or more sequencing reads with the same TSS contained different TESs, they were considered independent ROIs. Similarly, ROIs with different PAT length were regarded as independent, and we accepted those TSSs that have been described by others^20, 41–47^. Advantages of the ONT MinION sequencing technique over the PacBio platform include cost-effectiveness, higher read output, and the ability to read sequences within the range of 200 to 800bps, in which PacBio and the SRS techniques are less effective^48^. Although this method is hampered by high error rates, sequencing accuracy is not particularly detrimental to transcriptome analyses of well-annotated genomes, such as that of VACV^49^. In addition to the oligo(dT)-based 1D sequencing protocol from ONT, we prepared a random nucleotides-primed to detect the non-polyA(+) RNA fractions and to validate TSSs. A Cap-Seq approach was used to identify TSS positions. Together, these nanopore sequencing methods yielded about 535,000 viral sequencing reads. VACV ORFs are relatively short (**S2 Table**) and many overlap with each other. Because the PacBio MagBead loading protocol selects DNA fragments of less than 1kb^50^, potential monocistronic transcripts with a short ORFs are underrepresented or missing in these datasets. Nonetheless, in analyses of very long polycistronic and complex transcript, PacBio has better sequencing precision than ONT. Thus, combined use of these LRS methods eliminates the shortcomings of both. Native RNA sequencing can circumvent spurious transcription reads that are generated in PCR and RT reactions by false priming, template switching, or second-strand synthesis due to the lack of these techniques. The major disadvantage of dRNA sequencing is that a few bases are always missing from the 5’-ends of transcripts, and in many cases also from the 3’-ends also^51^. Under these sequencing conditions, it is assumed that the missing 5’-end nucleotides are the consequence of perturbed base calling due to premature release of the mRNA from motor protein, which hasten progress of RNA molecules across pores. The frequent absence of PATs is explained by the miscalling of adapter nucleotides that are ligated downstream of the PAT, thus muddling of the raw signal of the downstream ‘A’ homopolymer. The present dRNA-Seq analysis returned a total of approximately 195,000 reads.

**Fig 1.**
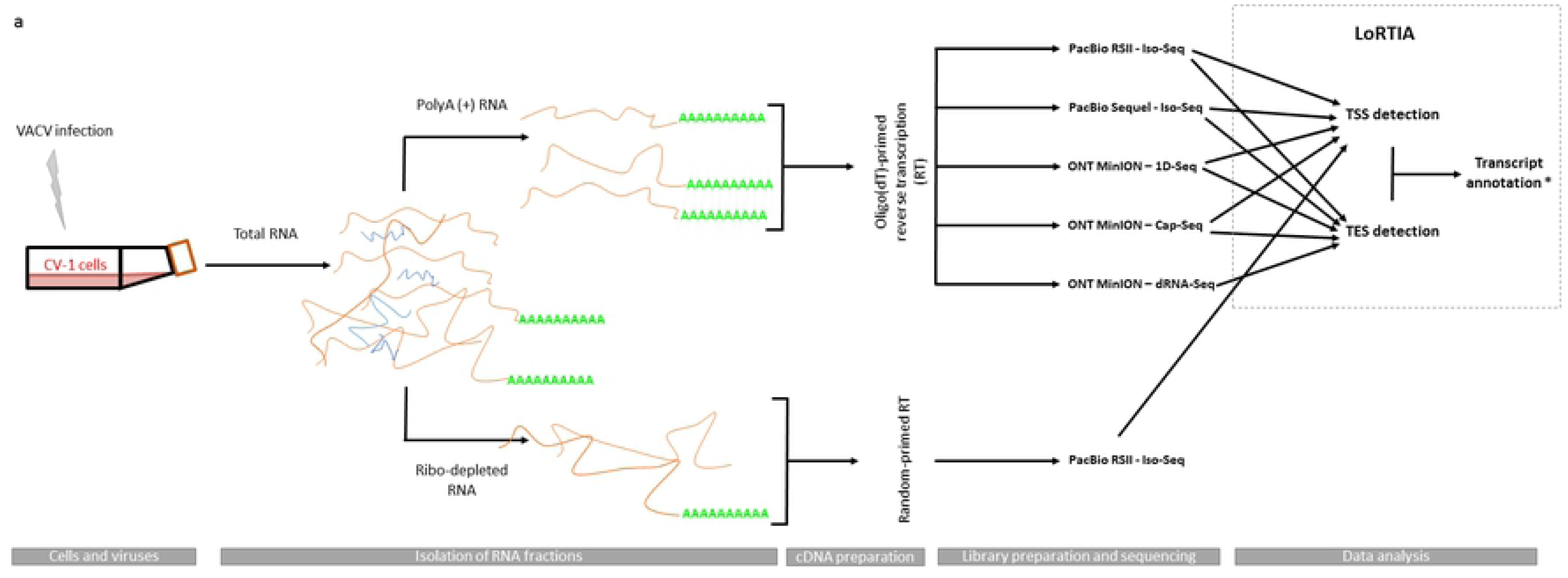
**Draft workflow of the study;** infection, sequencing, and data integration strategy for transcript annotation

This study demonstrates an extremely complex transcription pattern along the entire viral genome. Our sequencing approaches detected transcriptional activity at every nucleotide of the VACV genome (**Fig 2**). Numbers of annotated ORFs vary between 201-263^4, 52^ depending on the strain and study. Accordingly, we identified 218 ORFs in the WR strain (GenBank accession number is LT966077.1) using *in silico methods*, and 184 of these were expressed alone on separate RNA molecules. We detected 8,191 unique putative transcripts using the LoRTIA pipeline (**S3 Table A**). Under the strict criteria of detection by two different techniques and/or in three independent experiments, this number decreased to 1,480 transcripts (**S3 Table B**), including potential mRNAs, non-coding transcripts, or RNA isoforms. Transcripts annotated by LoRTIA represented 175 ORFs (**S3 Table B and C**). Our dataset contains full-length transcripts for about 20 additional ORFs (**S3 Table D**), but no full-length reads were detected for about 25 ORFs. These regions contain several transcriptional reads without precise TSS and/or TES positions. Specifically, starts or ends were within annotated ORFs (e.g.: O1L) or on complementary DNA strand (e.g.: A11R region). Our data show an enormously high level of read-through transcription. Our analysis detected ‘regular’ (**S1 Figure**) and ‘chaotic’ (**Fig 3**) regions of the viral genome. At the ‘regular’ genomic segments viral genes produce transcript with more or less precise TSSs and TESs, whereas at the ‘chaotic’ regions (such as the genomic loci of O1L, O2L, I1L, I2L, I3L, A10L, A11R, A12L, A18R genes, or the genes within the region of L genes) a large amount of TSSs and TESs diversity are produced. Some VACV gene clusters exhibited a genomic organization that resembled that of herpesviruses, with tandem genes producing overlapping 3’co-terminal RNA molecules.

**Fig 2.**
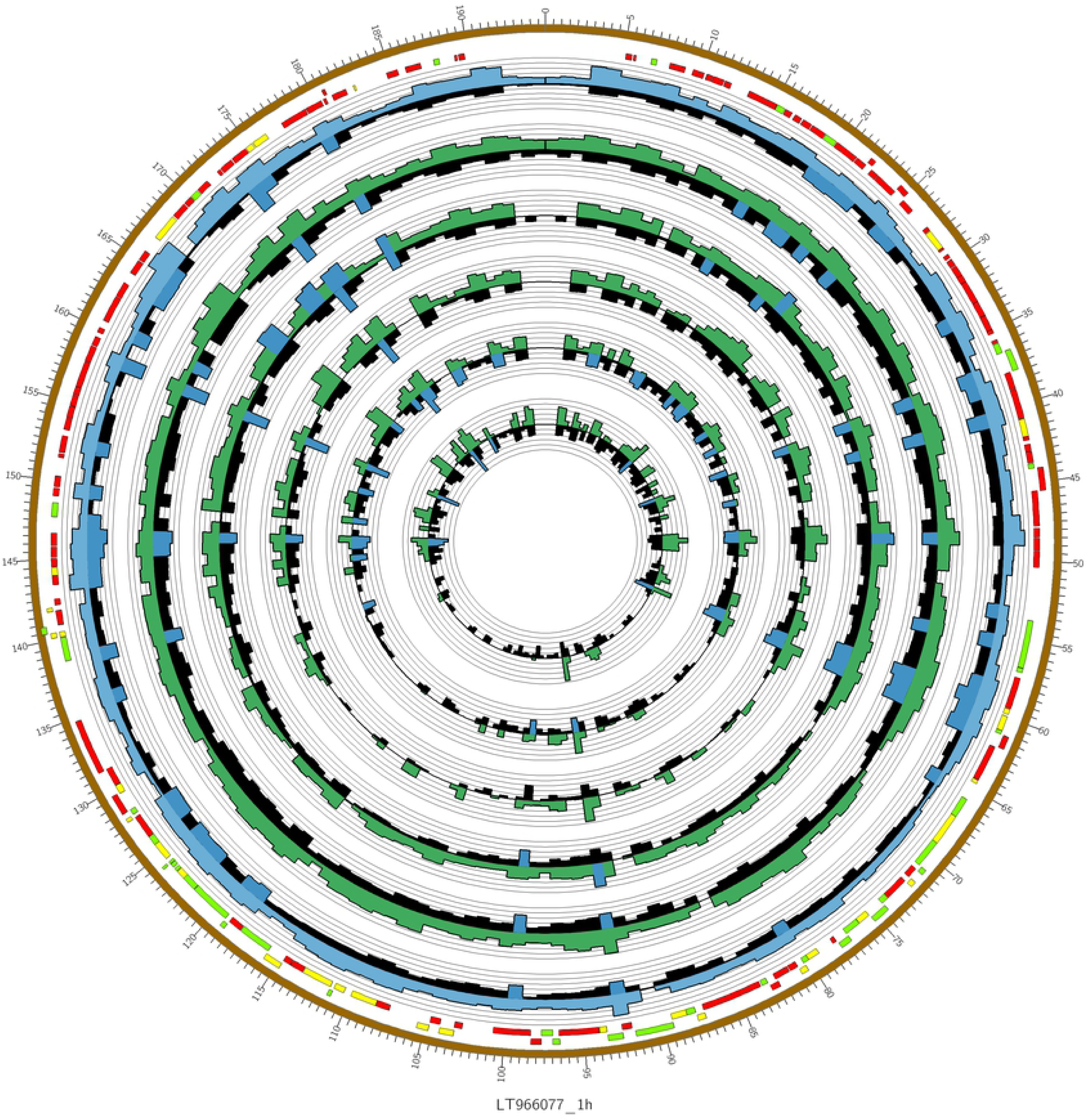
**Depths of VACV read coverage across the viral life cycle;** transcripts were obtained at six time points and were sequenced using PacBio Sequel (inner radius) and Oxford nanopore technology (ONT) MinION (outer radius) platforms and were visualized in a Circos plot as concentric circles; each circle represents an individual time point with early to later time points represented as inner to outer circles, respectively.

**Fig 3.**
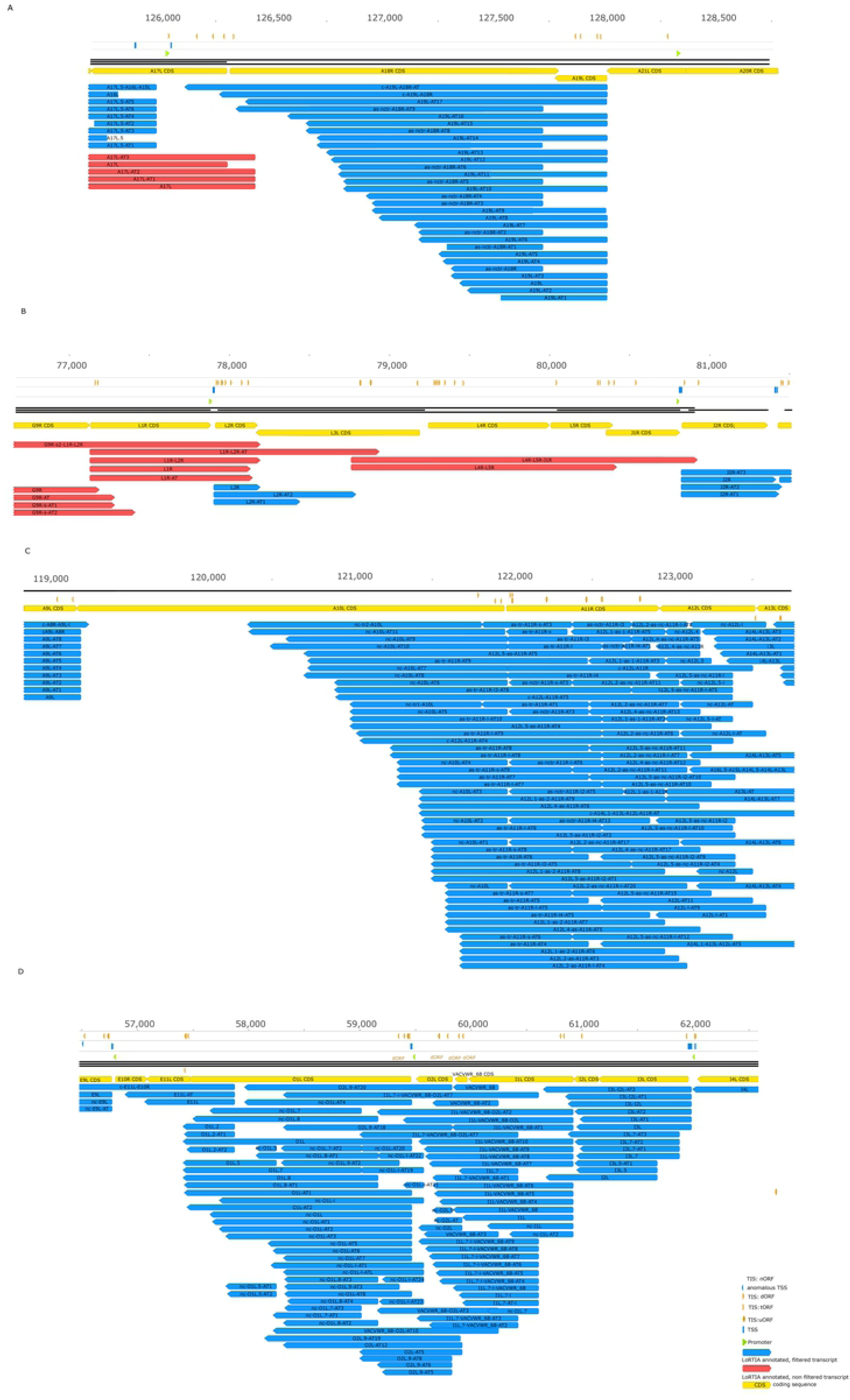
**Complexity of VACV ‘chaotic regions’; a.** high expression of antisense transcripts within the A18R locus; **b.** transcription start and end sites within the open reading frames (ORFs) of L genes; **c.** extremely high expression levels of antisense RNAs and other non-coding transcripts at the A10L-A12L region; **d.** enormous numbers of non-coding transcripts and transcript isoforms within the O1L– I3L genomic segment. All transcripts were identified using the LoRTIA toolkit.

### Novel putative protein-coding genes

Herein, we identified novel genes by identifying (a) new genomic positions, (b) embedded genes (5’-truncted in-frame ORFs within larger canonical ORFs), and (c) short upstream ORFs (uORFs, preceding the main ORFs). We considered the most abundant transcript isoform specified by a given gene as the canonical (main) transcript. (**S1 Note**). The novel ORFs and transcripts were named according to the common (Copenhagen) HindIII fragment letter/number-based nomenclature^46^. All VACV transcripts (identified by the LoRTIA pipeline) are presented in a Geneious file available at **figshare** (details: **S2 Note)**.

**(a) Genes at novel genomic** positions We detected two novel putative mRNAs that are encoded in intergenic genomic locations (B25.5R/C19.5L, C9.5L) between two divergently oriented genes and have short ORFs (both are 51bps). The TSS positions of these transcripts have been reported in previous publications^15, 20^ (**S4 Table**). Our analysis showed that these ORFs are expressed alone in separate transcripts and/or as upstream ORFs (uORFs) located 5’ of canonical ORFs in longer transcripts.

**(b) Embedded genes** A characteristic feature of the VACV genome is that it contains several shorter in-frame ORFs, and these have been shown to be translated^13, 41^. Full-length sequencing identified 49 novel putative embedded genes with 5’-truncated in-frame ORFs, and these were shorter than *in silico* annotated canonical ORFs (**Fig 4**). These putative genes specify more than 300 transcript isoforms. We assumed that the TISs of these internal ORFs were the closest in-frame ATGs to canonical host ORFs. A large number of TISs were detected previously using ribosome profiling^13^. For these, we identified 22 unique promoters using our in-house scripts, but could detect only six transcripts containing ORFs (**S5 Table**). Theoretically, these novel transcripts follow either alternative transcription initiation from distinct promoters or by leaky transcriptional initiation.

**Fig 4.**
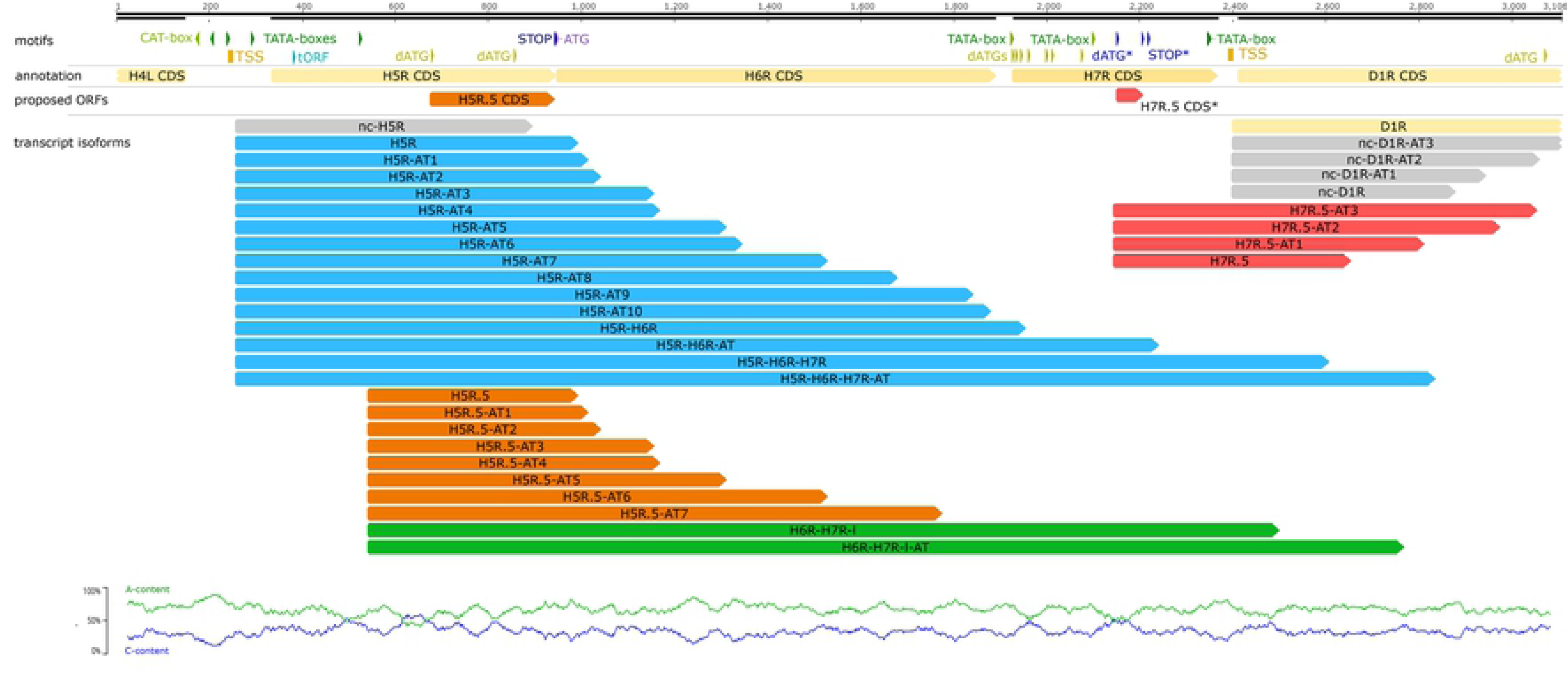
**Schematic of two examples of the identified putative embedded genes;** we detected a number of 5’-truncated transcripts containing short in-frame ORFs. This figure shows transcripts from H5R.5 and H7R.5 ORF regions.

**(c) Upstream ORFs** Upstream ORFs (uORFs) are considered common regulatory RNA elements in mammalian transcripts^53^, and 5’-UTR of uORF-containing transcripts are reportedly ≥ 75 bp in length^53^. Several *in silico* and experimental studies show that 40–50% of human and mouse mRNAs contain at least one uORF^54–55^. We applied strict criteria for filtering data obtained by the LoRTIA toolkit and revealed that 25 previously annotated VACV genes express longer TSS variants than the canonical transcripts. On average, 5’-UTR of the main transcript isoforms were 101 bp (**S2 Figure**), whereas the longer variants were 258 bp (**S6 Table**). Most of these transcript isoforms (64%) contain at least one ORF preceding the main ORF with an average 5’-UTR of 331 nts. We set the minimal size limit of these uORFs to 9 nts comprising at least one triplet between the start and stop codons, as described by Scholz and colleagues^56^. We also set a maximal ORF length of 90 nts and considered transcripts with longer upstream ORFs as bicistronic. Eleven transcripts with 5’-UTR variants were contained multiple uORFs. G5.5R was the only gene in which the longer transcript isoform contained a single uORF. The VACV transcriptome is distinguished by high number of TSS and TES positions and combinations per ORF. The same TSS can be used by the longer variants of downstream genes and by a 5’-truncated, potentially protein coding transcript of the upstream gene.

In many such cases, the overlapping part of the two transcripts (the long 5’-UTR of the downstream gene and the 5’-truncated version of the upstream gene) contains one or more ORFs that can be considered uORF (within the L variant) and/or a novel potential protein-coding genes, as for A1L, A34R, and H3L regions, and C5L). The present novel transcripts of possible uORFs are listed in **S6 Table.**

### Non-coding transcripts

In the LoRTIA analyses presented in **S3 Table B**, we identified 356 novel putative long non-coding RNAs (lncRNAs) of longer than 200 bps. We also identified 13 potential short ncRNAs (sncRNAs) with lengths varying between 121 and 193 bps.

#### Antisense RNAs

With few exceptions, we detected transcriptional activity from both DNA strands along the entire viral genome, and mRNAs and antisense (as)RNAs expression levels varied between genomic locations. Relative low asRNA expression was detected in almost every ORFs. After applying strict criteria to the LoRTIA tool, 100 antisense transcripts were located at the genomic loci A5R (a single transcript), A11R (89 transcripts), and A18R (10 transcripts) (**Fig 5**).The A11R antisense transcripts are various combinations of six TSS and 40 TES positions, whereas antisense A18R transcripts have the same TSSs but varying termination sites. Some of these transcripts contain small ORFs that may be coding RNAs (**S2 Note**). Practically, the entire VACV genome express asRNAs, but in most cases in a very low abundance, that is why the LoRTIA tool was unable to identify these molecules.

**Fig 5.**
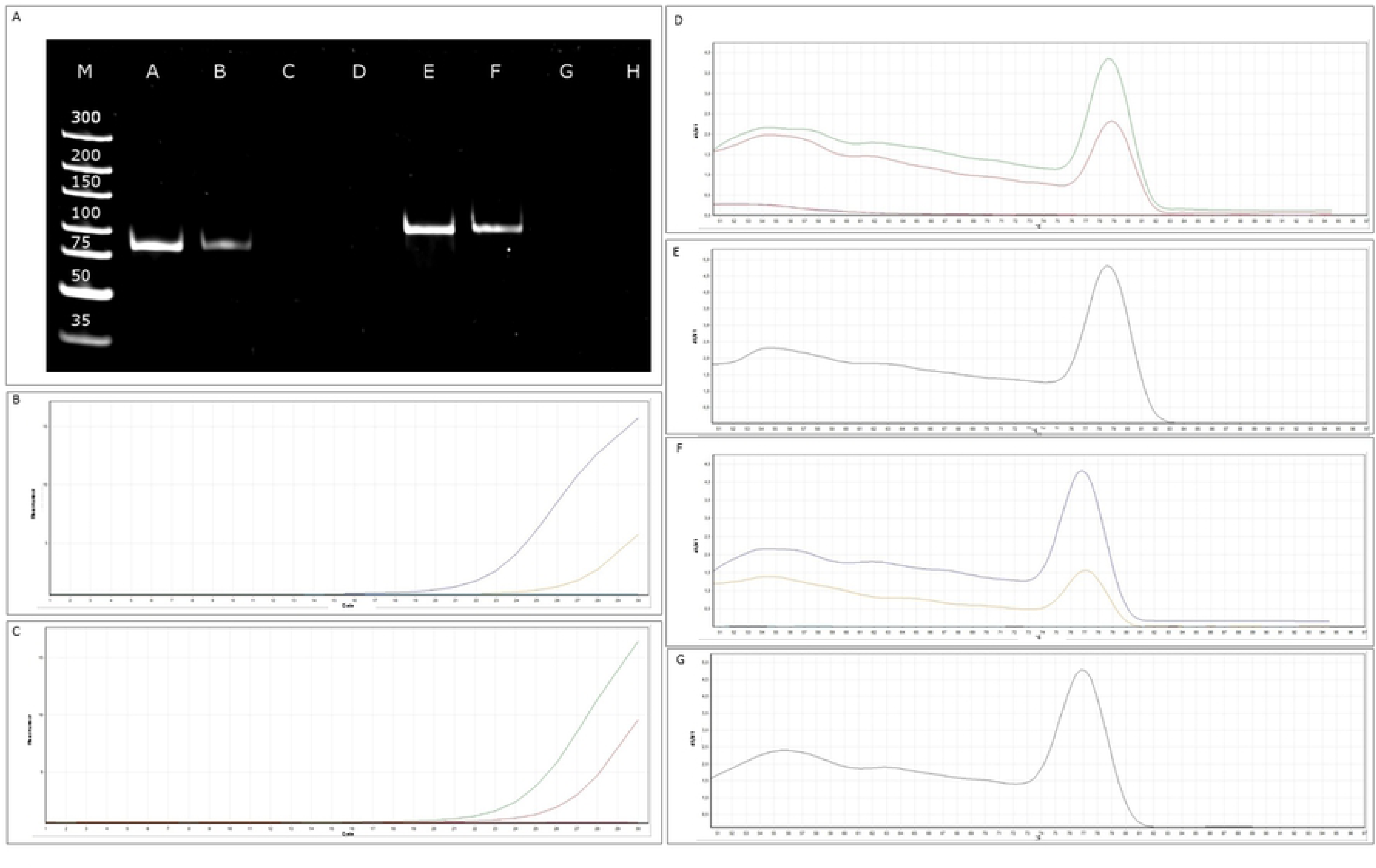
**Detection of antisense RNA expression from the complementary DNA strands of two VACV genes using qRT-PCR. A**. polymerase chain reaction (PCR) products of the A11R transcript and its antisense partner. **Abbreviations**: M: molecular weight marker; A: A11R antisense RNA; B: A11R mRNA; C: A11R antisense RNA – NO-RT control; D: A11R mRNA – NO-RT control; E: A18R antisense RNA F: A18R mRNA; G: A18R antisense RNA – NO-RT control; H: A18R mRNA – NO-RT control. **B**. amplification curves of the *A18R* gene and its antisense transcripts. **C**. Amplification curves of the A18R gene and its antisense transcripts. **D-G.** Melting curves are shown to demonstrate the specificity of the amplification. The curves of transcripts cross the threshold line within 19.5-23.4 cycles, whereas the curves of NO-RT controls remain flat. **D**. A11R mRNA and antisense RNA. **E**. A11R DNA control. **F**. A18R mRNA and antisense RNA **G**. A18R DNA control.

### Intragenic non-coding transcripts

After filtering with strict criteria, we identified 260 3’-truncated transcripts from the LoRTIA dataset and considered these non-coding transcripts because they lacked in-frame stop codons. These transcripts were found within 41 viral genes and 36.15% of them were located within the O1L region. The A37R (8.85%) and A10L (5.38%) regions and the F12L-F13L gene cluster (11.15%) also contained substantial numbers of 3’-truncated RNAs, and cumulatively, 62% of the 3’-truncated ncRNAs were located within five ORFs (**S3 Table B**).

#### Replication-associated transcripts

The VACV genome contains multiple replication origins^57^ (Oris) located near the genomic termini. Oris are overlapped by many non-coding transcripts that are expressed in low abundance, but the TSSs and TESs of these ncRNAs were not identified by LoRTIA or other methods because the transcripts vary widely in size. Nonetheless, we demonstrate that these RNAs are expressed from 4 h post infection (p.i.), and they may play similar roles as herpesvirus replication-associated RNAs^58^.

### Novel mono-, bi- and polycistronic RNAs

Since SRS fails to detect full-length transcripts, every RNA molecule identified by LRS techniques can be considered to be novel. At the ‘regular’ genomic regions we detected 135 monocistronic coding transcripts of previously annotated ORFs using the LoRTIA software suit, and 12 additional mRNAs did not meet the LoRTIA criteria due to their low abundance (**S7 Table**). Similar to other viruses and prokaryotes, but unlike the eukaryotic organisms, VACV reportedly express bi- and polycistronic transcripts^59–61^. Yet although all genes are translated from prokaryotic polycistronic mRNAs, experimental evidence is lacking for translation of downstream genes from polycistronic mRNAs of VACV. We detected 43 bicistronic, 137 tricistronic, 92 tetracistronic, 15 pentacistronic, and 5 hexacistronic RNA molecules (**S3 Table B and C**). Yet not all of these 294 VACV polycistronic transcripts are ‘regular’ because many are initiated or terminated within the ORFs.

### Complex transcripts

Herein, complex transcripts are those RNAs containing at least one anti-polar ORF on multigenic transcripts. Using LoRTIA, we detected 30 complex RNA (cxRNA) molecules (**S8 Table, S3 Table B, C and D, S2 Note**) localized within nine genomic segments. We considered six of these as TES isoforms of the A12L–A11R complex transcript, five cxRNAs as TSS and TES combinations within the A14L-A13L-A12L-A11R region, and four transcripts as alternative TES variants of VACVWR_161-A38L RNA. Three additional complex transcripts were excluded in LoRTIA analyses due to low coverage at this genomic region. Because VACV genes stand in tandem orientation with each other, few convergently and divergently positioned gene pairs are present, leading to few cxRNAs compared with those in herpesviruses. Two cxRNAs (c-A21L-A20R, c-G3L-G2R) were considered non-coding because their upstream genes were antisense on the transcript. The remaining complex transcripts were categorized as mRNAs, because their first ORFs were in the sense direction. We detected cxRNAs with convergently-oriented genes at E10R-E11L and A8R-A9L genomic regions. Fully overlapping convergent cxRNAs were also detected in the E10R-E11L region (**S3 Figure**).

### LRS reveals highly complex isoform heterogeneity

Transcript isoforms include 5’- and 3’-UTR length variants of RNA molecules. Using RNA-Seq tag-based methods with SOLiD and Illumina sequencing platforms, Yang and co-workers determined 5’- and 3’-ends of VACV E mRNAs with precision^15, 20^ and mapped terminal sequences of I and L transcripts^41^. These investigators also determined expression profiles of I and L RNA-products^16^. These studies identified several hundreds of TSSs and PASs, but uncertainty of 3’-ends of RNAs remained due to the high complexity of gene expression and overall read-throughs of upstream genes at late time points of infection^15^. We annotated 1,073 TSSs of VACV transcripts, 987 of these have not been previously detected (**S9 Table**). Seventy percent of the present TSSs were located within ± 10-nt intervals, whereas 93% of the described TSSs occurred in ± 30-nt intervals and 18% of TSSs were matched with base-pair precision to those described by Yang and colleagues^15, 16, 20, 41^ (**Fig 6A**). Altogether 898 TSSs were detected in the ‘regular’ regions. The previously published TSS positions are depicted in **figshare (S2 Note)**. TESs of early transcripts were increasingly heterogenic at late time points of infection, potentially because transcription termination signal recognition depends on different factors in early- and PR phases^13, 15–16, 20^. Because viral TESs were highly variant (**S4 Figure**), we analyzed positions within a ± 30-nt broad scale and found that 24% of our annotated TESs matched the positions of the polyadenylation sites described by Yang *et al*.^16^ (**Fig 6B**). Moreover, most genes express multiple transcript isoforms (median = 4, mean = 6.95). In the extreme cases of N1L, O2L, I1L, and A12L regions, more than 30 transcript isoforms were expressed (**S3 Table B**; **S5 Figure**). Moreover, the TSS positions of about 30 annotated ORFs within the VACV genome, particularly in the A8R-A17L region, could not be annotated in ours or previous studies (**S3 Note**). Yang and colleagues described a 15-nucleotide consensus promoter core motif upstream of E genes, but these sequences were also present at other genomic locations. We identified this motif at nine novel positions upstream (C9.5L, F4L, F7L, J3R, A15R, VACWR_161, and A51R) and inside (A36R and VACWR_169) ORFs. Cis-regulatory elements have not previously been annotated within these regions (**S2 Note**). However, these core motifs are not present at the ‘chaotic’ genomic regions. However, these core motifs are not present in the chaotic genomic regions and most novel transcripts had TESs located about 50 nucleotides downstream of UUUUUNU termination signals, as described by Yang et al^20^. The presence of several TESs that were not preceded by this motif suggests an alternative mechanism for polyadenylation-site selection. TSSs with TESs can only be matched by LRS techniques, which are therefore crucial to analyses of the complexities of the VACV transcriptome. In chaotic genomic segments, almost all base positions served as TESs within relative long sequences. Transcription initiation was also stochastic, especially at these regions. Accordingly, our analysis revealed that VACV genes express transcripts with multiple shared TSS (**Fig 3A**, **B, C** and **S4 Fig A**) and TES positions (**Fig 3C** and **D, S3 Figure B**). Distributions of TSSs and TESs along the VACV genome are presented in **S6 Figure.** We identified TSSs for 90% of the promoters described by Yang *et al.*, (2010), with exceptions of G2R, J6R, A18R, A20R, and five hypothetical genes located in the repeat region of the VACV genome. We used *in silico* promoter predictions to detect potential TATA- and CAAT-box consensus sequences of the present TSSs. We analyzed 146 TSSs without previously annotated promoters and found TATA-boxes in 88 (**Fig 7, S10 Table**), and CAAT-boxes in 5 of these (**S10 Table**). The median distance between TSS and putative TATA-box was 54 nts. Yet, no significant differences in promoter–TSS distances were identified between 5’-truncated and canonical transcript isoforms, suggesting that cis-regulatory structures were similar in the two types of VACV genes (**Fig 7**).

**Fig 6.**
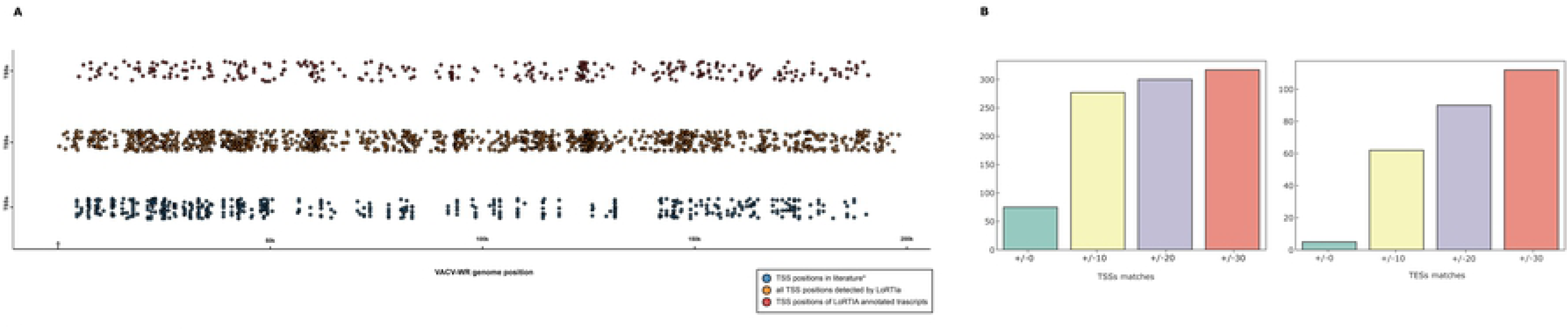
**A Jitter plot visualization of transcriptional start site distribution across the VACV genome;** distribution of TSSs of our annotated transcripts are shown at the top (red dots); the TSSs detected using LoRTIA are depicted in the middle (orange). TSS positions described by others^15–16, 20, 41^ are located at the bottom of the figure (blue). **B.** TSS and TES positions detected by LoRTIA share the TSS and TES positions obtained from other studies (**S9 Table**) with different frequency. Bar charts represent the fall of TSS and TES positions within a +/-10-20-30 sliding window.

**Fig 7.**
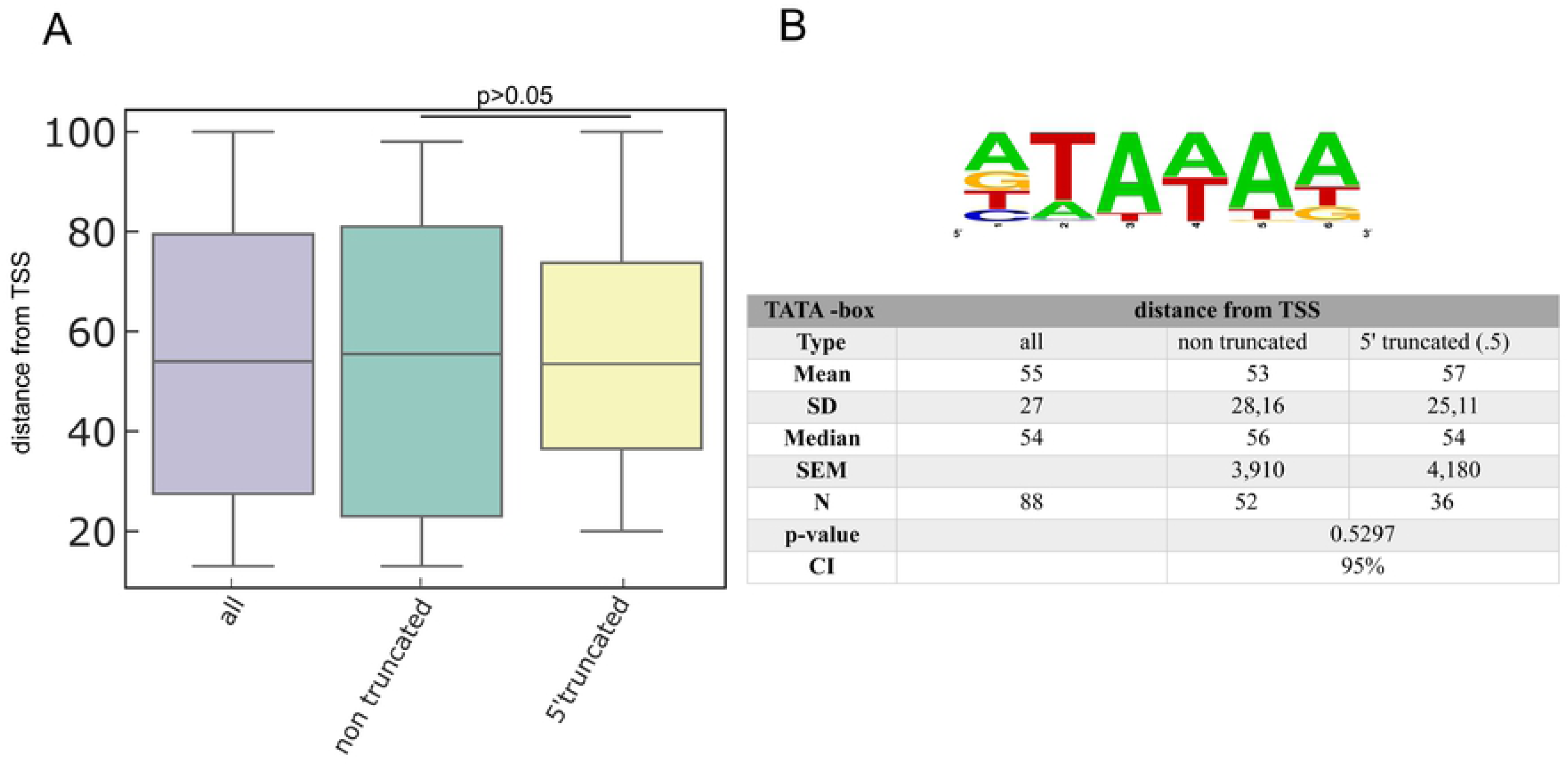
***In silico* analysis of promoter elements; a.** the bar chart shows average distances between the TATA boxes and TSSs. **b.** The WebLogo shows the consensus sequence of the identified TATA boxes.

### Transcriptional overlaps

Using multiplatform analyses, we revealed an extremely complex meshwork of transcriptional overlaps, which were expressed by all viral genes. Cumulatively, 154 tandem, 32 divergent, and 32 convergent gene pairs were found in the VACV genome, with 21 overlapping ORFs between tandem genes, 13 between convergent gene pairs, and 5 between divergent gene pairs. Moreover, closely-spaced tandem genes can form partial and full parallel overlaps. We also detected 106 parallel transcriptional overlaps between tandem genes (**S11 Table**), reflecting overlap by canonical transcripts, and by longer TES or TSS isoforms of upstream or downstream genes, respectively. Seventeen of the 32 convergent gene pairs formed UTR-UTR overlaps (**S11 Table**), whereas only 15 gene pairs formed UTR-ORF overlaps. Twelve of 16 divergently-oriented gene pairs were also found to form transcriptional overlaps with each other (**S11 Table**). Polycistronic and complex transcripts are also consequences of transcriptional overlaps (**S12 Table**).

### The dynamic transcriptome of VACV

#### Transcript kinetics

As a major advantage, LRS methods perform end-to-end sequencing of entire RNA or cDNA molecules, allowing identification and kinetic characterization of transcript isoforms. We performed kinetic analyses of LoRTIA transcripts using hierarchical clustering and identified five distinct clusters in the dataset (**Fig 8**). K-means clustering was used to describe the temporal expression levels of these clusters (**S7 Figure**), and following expression profiles were obtained: (1) the *early-down* group had continuously decreasing Rt values (see Methods) from the start of infection; (2) the *mid up-down* cluster had a local maximum between 2 and 3h p.i.; (3) the *late up-down* cluster of transcripts reached maximum of Rt values at 6h p.i.; (4) the *late up* cluster of transcripts had a clear late activity reaching maximum Rt values at 8h p.i., and (5) most viral transcripts exhibited minor changes throughout the viral life cycle (*constant* cluster) (**S13 Table**). The dynamics of various transcript categories are presented **Figs 9** and **10**, which show TSS and TES isoforms, polycistronic RNAs and Ori-associated transcripts. Many RNA isoforms and mono- and polycistronic transcripts are differentially expressed throughout the viral life cycle.

**Fig 8.**
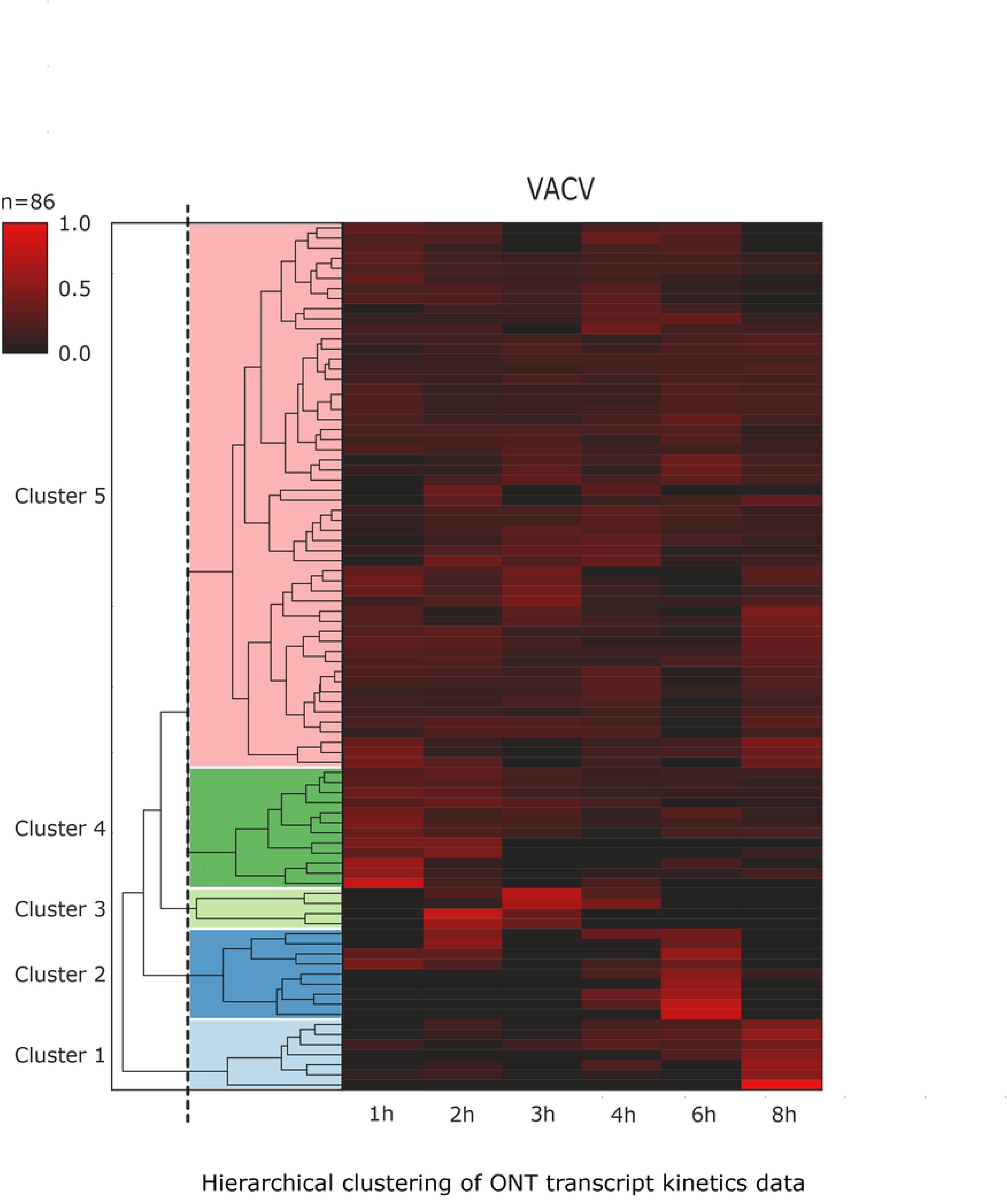
**Expression profiles of the most abundant dynamic vaccinia virus (VACV) transcripts and transcript isoforms represented by a heatmap matrix;** Rt values of the viral transcripts show distinct expression profiles in the five kinetic clusters. Each row represents changes in relative expression levels of a RNA. Red rectangles indicate high relative expression values and black rectangles indicate low relative expression values. All transcripts were identified using the LoRTIA pipeline.

**Fig 9.**
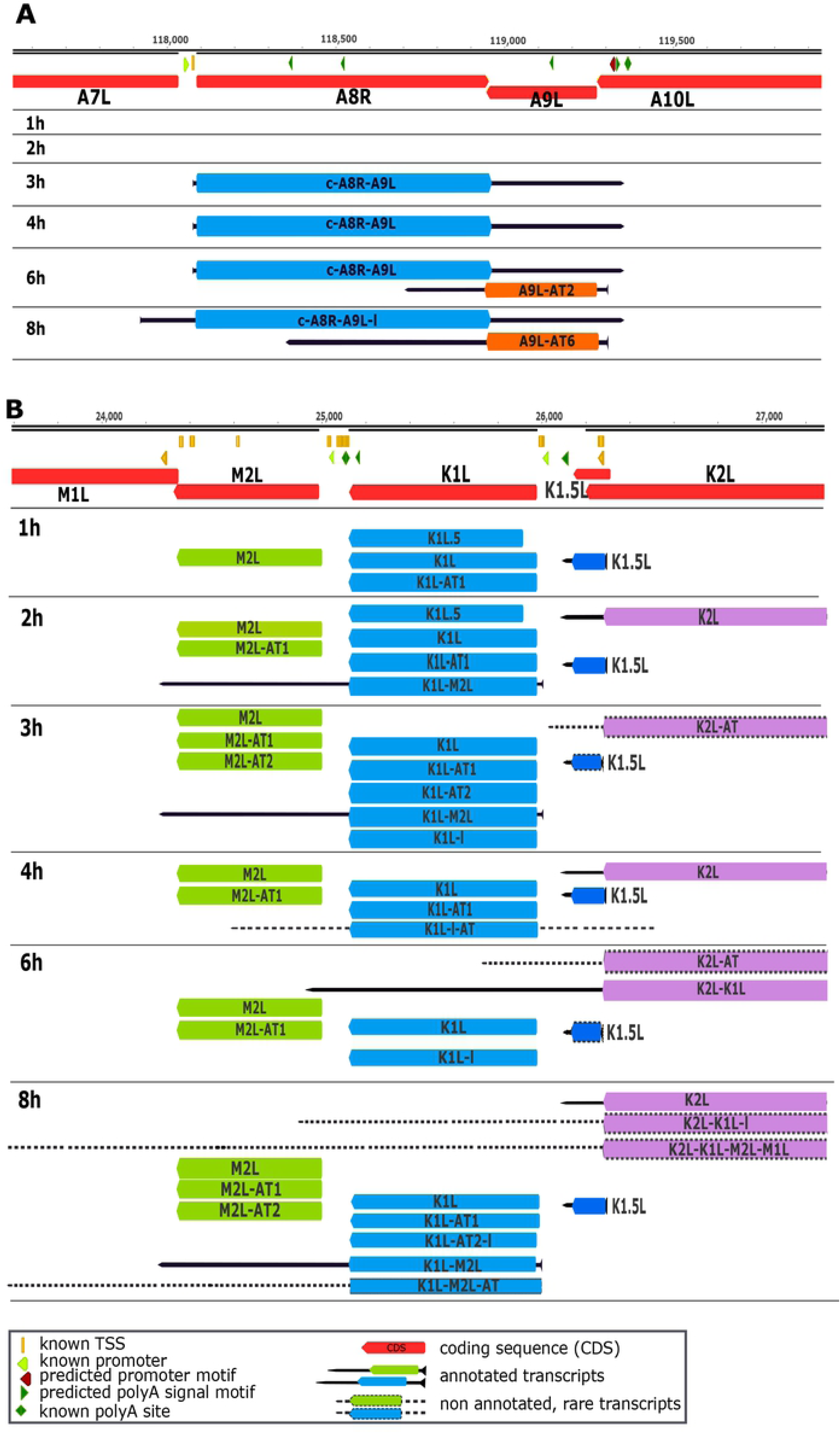
**Dynamic VACV transcriptome; A.** time-dependent changes in quantities of TSS isoforms of the A8R-A9L complex transcript and the most abundant AT RNA isoforms of the *A9L* gene; **B.** expression levels of various transcripts and transcript isoforms within the M1L-K2L region, including transcriptional end site (TES) isoforms, mono- and bicistronic variants, and novel putative protein-coding genes; transcript structures and counts were determined using the LoRTIA software suite.

**Fig 10.**
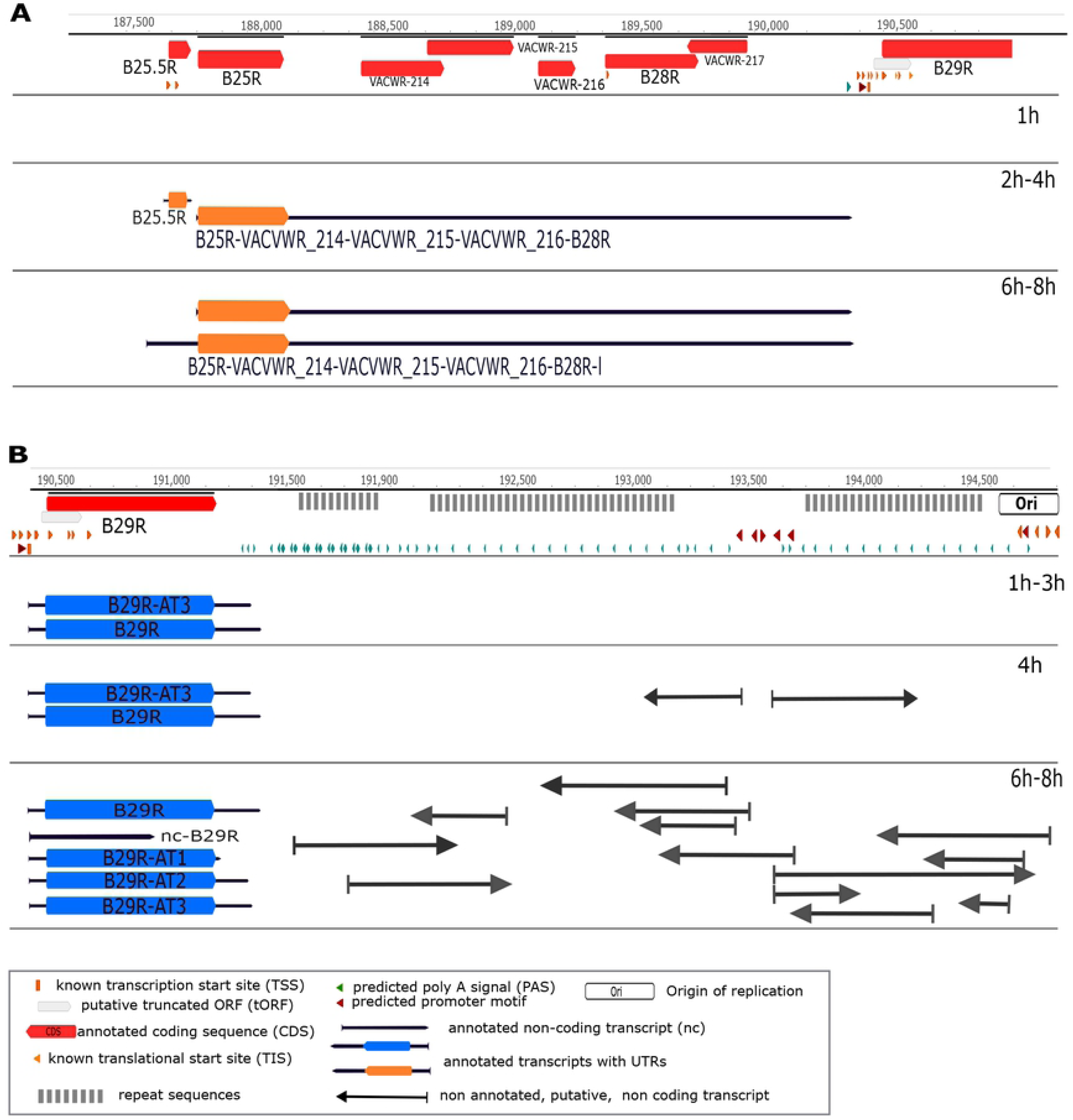
**Expression dynamics VACV transcripts A.** Dynamic change of mono-, bi-, and polycistronic transcripts. **B**. Expression dynamics of the replication-associated RNAs.

#### Dynamic expression of VACV ORFs

Hierarchical clustering according to ORF kinetics yielded five temporal clusters (**S8 Figure**). We summed transcripts with given ORFs alone or in the most upstream position of a polycistronic transcript (downstream ORFs are untranslated). We summed transcripts with given ORFs alone or in the most upstream position of a polycistronic transcript (downstream ORFs are untranslated) and showed similar ORF kinetics to those of their canonical transcript isoforms. Hence, the relative proportions of other transcript isoforms were generally low relative to the main isoforms, and therefore had no significant effects on the expression dynamics of ORFs.

#### Dynamics of the transcriptional read-throughs

TES isoforms are produced by passing of at least one transcription termination signal by the RNA polymerase, yet temporal expression of TES variants differed between genes. For some genes, all transcript isoforms appeared at every time point (WACVR_15, H5R, and their isoforms), whereas certain isoforms of other genes were expressed at single (N1L and B19R) or multiple time points (WACVR_15, H5R and their isoforms), whereas in other genes certain isoforms are expressed in a single (N1L, B19R), or multiple time points (F14L, O1L) (**S9 Figure**). According to our strict criteria, the A37R gene exhibited the highest variation in 3’-UTR with seven alternative ends. With some exceptions (e.g., WACVR_15), the expression profiles of 3’-UTR variants differed from each other throughout the viral life cycle.

### The host cell transcriptome

We used PacBio Sequel and ONT MinION LRS platforms to analyze full-length isoforms of the CV-1 cell line preceding and during VACV infection. MinION cDNA sequencing yielded a total of 1,375,219 reads, and 964,775 of these were mapped to the host genome with an average mapped read length of 583 nts. Sequel sequencing analysis yielded 479,179 ROIs of which 439,330 mapped to the host genome with an average mapped read length of 1,368 nt. The library preparation steps that mitigate loading bias of PacBio sequencing worked well for sample capture in host cells, but not so well for shorter VACV transcripts (**S10 Figure**). In contrast, MinION sequencing had no preference for read length, resulting in a higher yield of short reads mapped to the virus and a more realistic change in host/virus read proportions at each p.i. time point (**Fig 11**). Using the LoRTIA toolkit, we annotated a total of 478 TSSs, 2,011 TESs (**S14 Table**) and 24,574 splice junctions (**S15 Table**), all of these were represented by at least 10 reads. Analyses of sequence regions upstream of TSSs revealed that 43 contain canonical CAATT boxes at a mean distance of 104.913 nt with a standard deviation (sd) of 15.306, 880 contained canonical GC boxes at a mean distance of 60.095 nt (sd = 33.374), and 80 contained canonical TATA boxes at a mean distance of 31.13 nt (sd = 2.966). Vo Ngoc and colleagues demonstrated that initiator elements surrounding TSSs of human genes are more likely to be present if the transcript lacks a TATA box upstream of its TSS^62^. In our analyses, the BBCABW (B = C/G/T, W = A/T) initiator consensus was more pronounced around TSSs of TATA-less genes than around those with TATA boxes upstream of their TSSs (**Fig 12**). Specifically, 1,849 TESs contain canonical upstream poly(A) signals at an average of 26.681 nt (sd = 9.340), whereas ±50-nt regions contain U-rich upstream elements and G/U-rich downstream elements (**Fig 13**). We annotated a total of 12,287 introns, and 12,215 of these contained canonical GT/AG splice junctions, whereas 65 had GC/AG, and 7 had AT/AC splice junctions. In determinations of transcript annotations using the LoRTIA pipeline, we included only transcripts with TSSs and TESs that were identified from at least two techniques and in three samples. The average transcript length for host cells was 693.661 nt (sd = 962.62), and no significant deviations were observed during infection. Lengths of 5’-UTRs deviated around a mean of 52.956 nt (sd = 75.903), whereas the 3’-UTRs had a mean length of 295.219 (sd = 431.480) (**S11 Figure**). Transcripts that were represented by ten or more reads were considered certain, and those with less than ten reads were excluded from further analysis. These assessments revealed a total of 758 transcript isoforms in greater than ten reads, 207 with TSSs and TESs in ±10-nt intervals of previously annotated transcripts, 692 length isoforms, and 66 alternatively spliced isoforms. In total, 239 mRNA length isoforms differed from previously annotated transcripts in their TSS positions (including those with TSSs downstream of respective TISs), 19 in their TES positions, and 56 in both. Together, 31 transcript isoforms were found with TSSs downstream of previously annotated TISs, and contained curtailed forms of ORFs. These RNAs may be non-coding or may code for N-terminally truncated forms of their canonical proteins. A total of 177 transcripts were annotated as non-coding. The present ncRNAs include either isoforms of previously annotated ribosomal RNAs (rRNAs), or are truncated mRNAs lacking ORFs. The transcript isoforms are listed in **S16 Table**.

**Fig 11.**
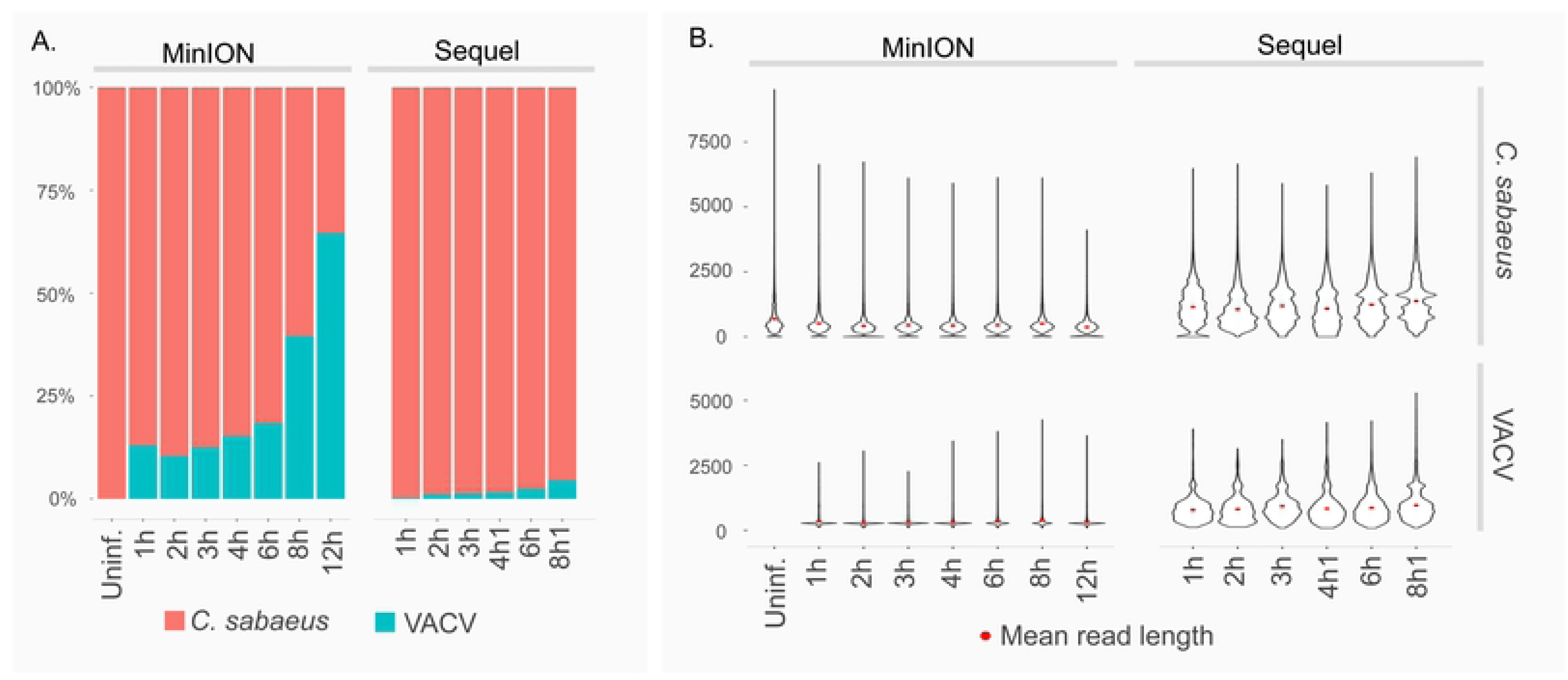
**Read counts and lengths of uninfected and infected samples at each time point**; **A.** fractions of reads mapped to host (*C. sabaeus*) and VACV genomes; dramatically reduced read counts of the host and an increased read counts of the virus are observable with viral life cycle progression in MinION data, whereas similar but much smaller changes were detected in the Sequel dataset. **B.** Mapped read lengths of the host and virus from MinION and Sequel sequencing from the uninfected sample and post infection (p.i.) samples; the panel shows the effect of library preparation for PacBio sequencing, yielding longer reads with greater abundance than those from MinION sequencing.

**Fig 12.**
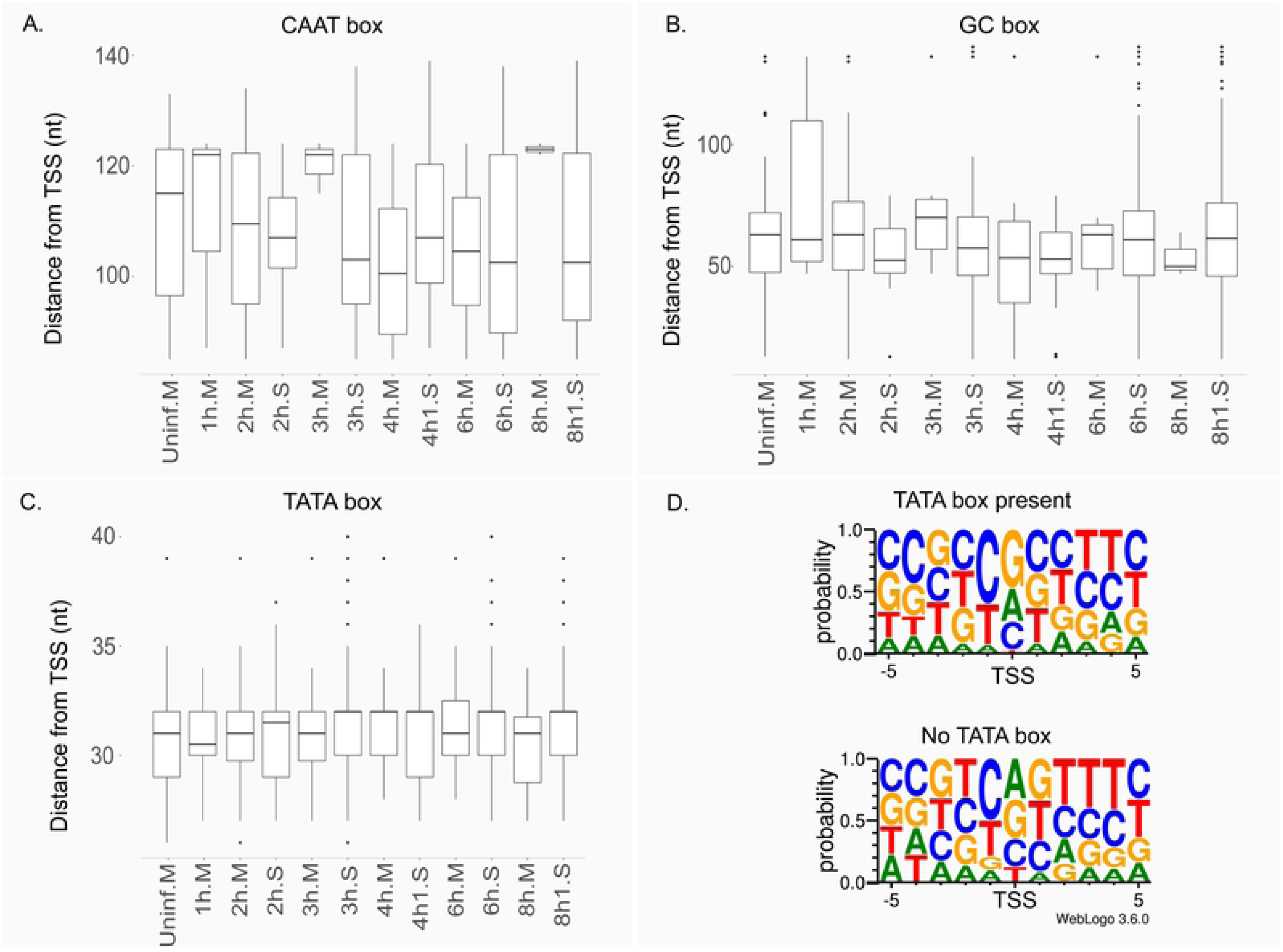
**The distance of the host promoter elements from the TSS or the transcription initiator sequence. A, B and C:** Letter ‘M’ following the sample name indicates MinION sequencing, whereas letter ‘S’ indicates Sequel sequencing. The horizontal lines in the box plots represent the median distance of the given sample. **D.** The transcriptional initiator region carries a BBCABW (B=C/G/T, W=A/T) initiator consensus sequence when no TATA box is present upstream of the TSS, whereas the consensus is missing in TSSs with a TATA box.

**Fig 13.**
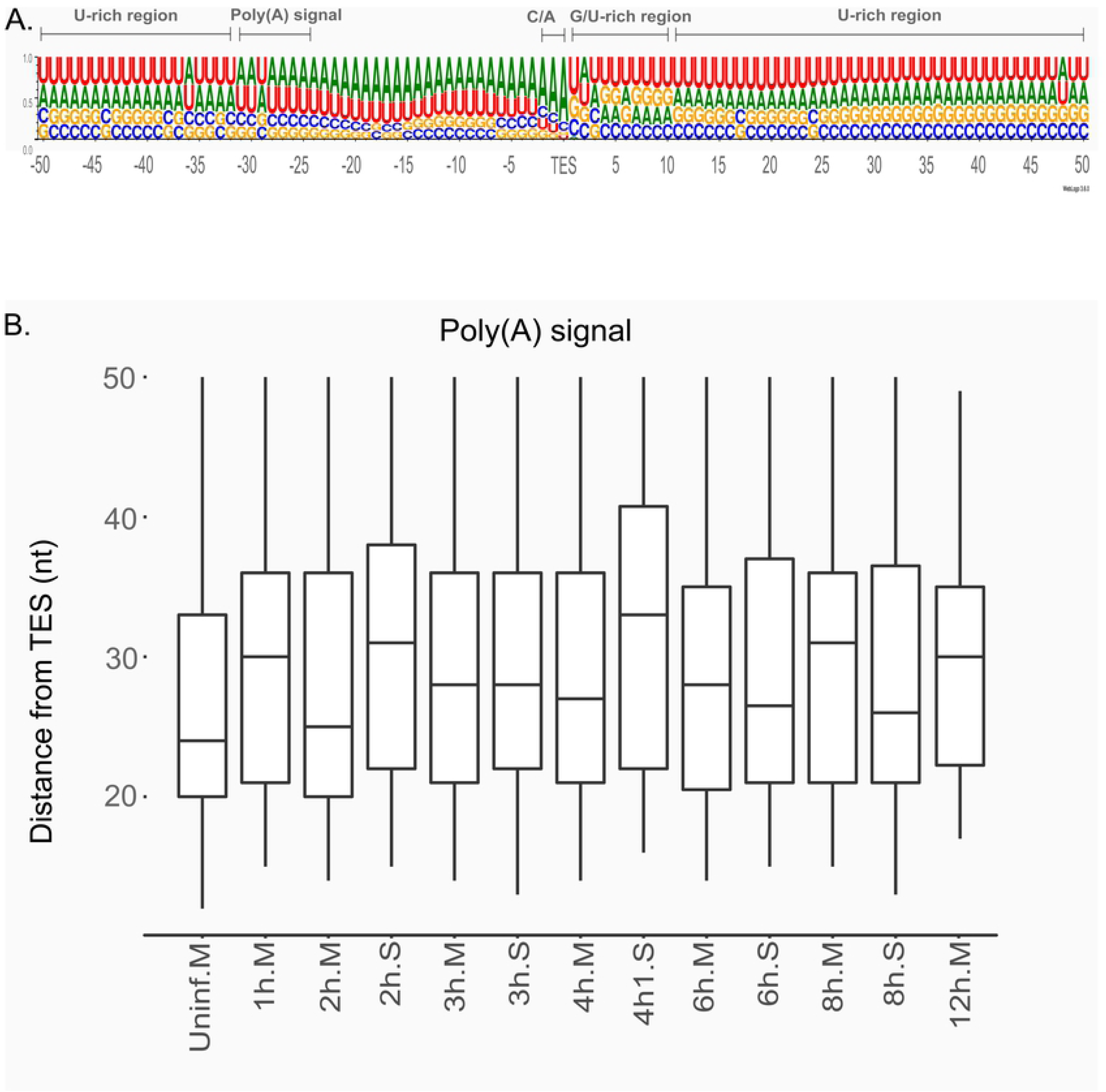
**The vicinity of the host TESs**. **A**. Nucleotide distribution surrounding the TESs of *Chlorocebus aethiops* were visualized using WebLogo showing canonical sequences signaling RNA cleavage and polyadenylation. B. The distance of the host’s polyadenylation signals from the TESs in the uninfected and the p.i. sample. The letter ‘M’ following the sample name indicates MinION sequencing, while letter ‘S’ Sequel sequencing. The horizontal lines in the box plots represent the median distance for the given sample. No significant change in the distance of TESs were observed during the viral infection. The TES positions were determined using the LoRTIA toolkit.

### Temporal response of the host transcription to the VACV infection

Using microarray technology, Rubins et al.^63^ and Bourquain et al.^64^ analyzed the impacts of VACV infection on host gene expression, but detected relatively few differentially expressed genes, potentially because this virus replicates in the cytoplasm. A recent temporal proteomic study also showed that VACV infection affects very few host genes^65^. In this study, we categorized five distinct clusters of 768 highly expressed host genes with respect to their responses to viral infection. Among (1) *early up* genes, no or very low expression levels were observed before virus infection, but consistently high expression was observed at all later sampling points. (2) *Early down* transcripts were highly expressed before virus expression and were then constantly absent or had low expression levels at all later time points. (3) *Early up/down* transcripts had no or low expression before virus infection, high expression from 1 hour after infection and no or low expression at later sampling points. (4) *Mid up* transcripts had no expression before virus infection and peaked and plateaued at 2 or 3 hours p.i. (5) *Constant* transcripts had no significant changes in relative expression levels over the course of our experiments (**Figs 14 and 15**).

**Fig 14.**
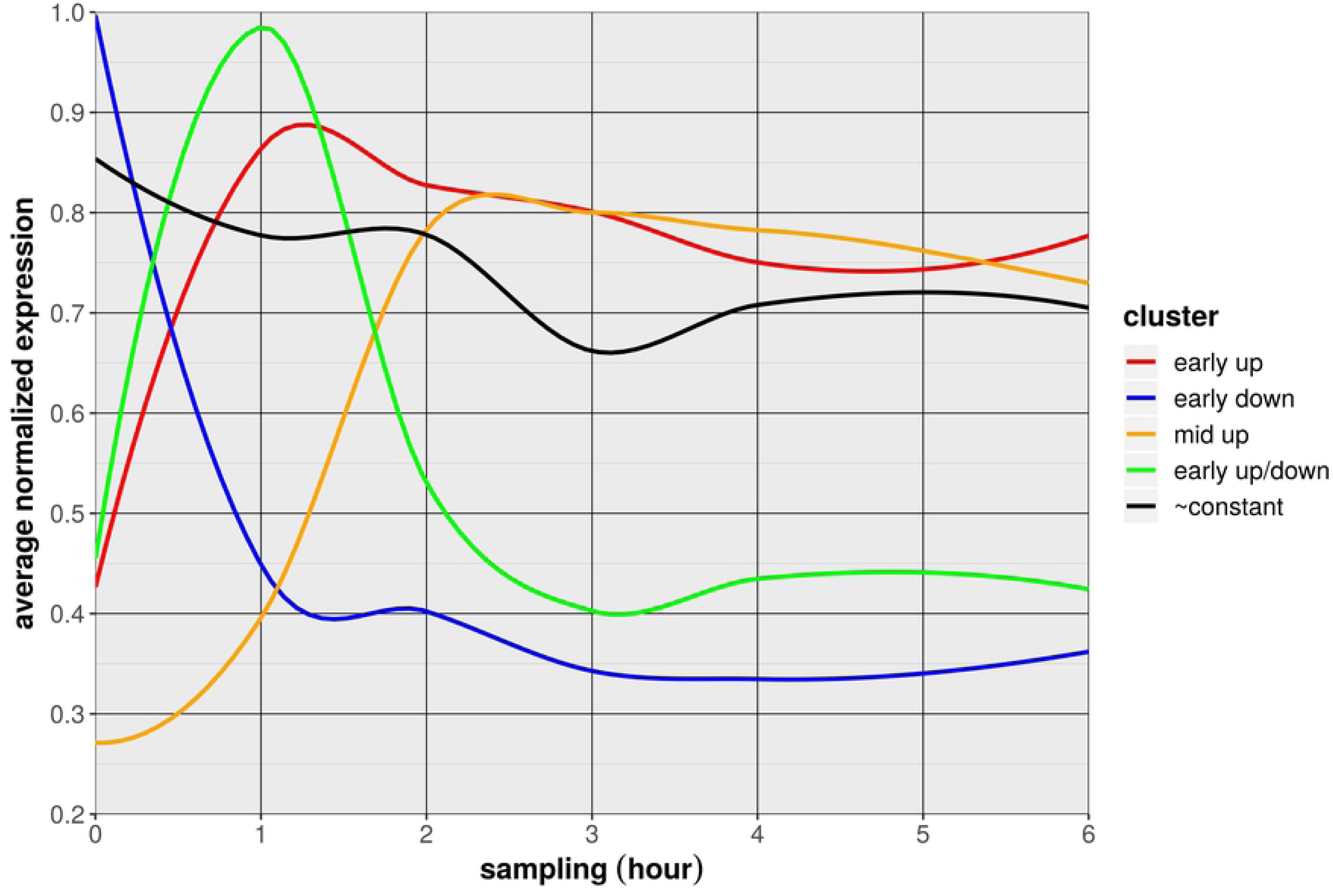
**Expression changes of the highly abundant host genes during viral infection.** The expression pattern of the host gene clusters.

**Fig 15.**
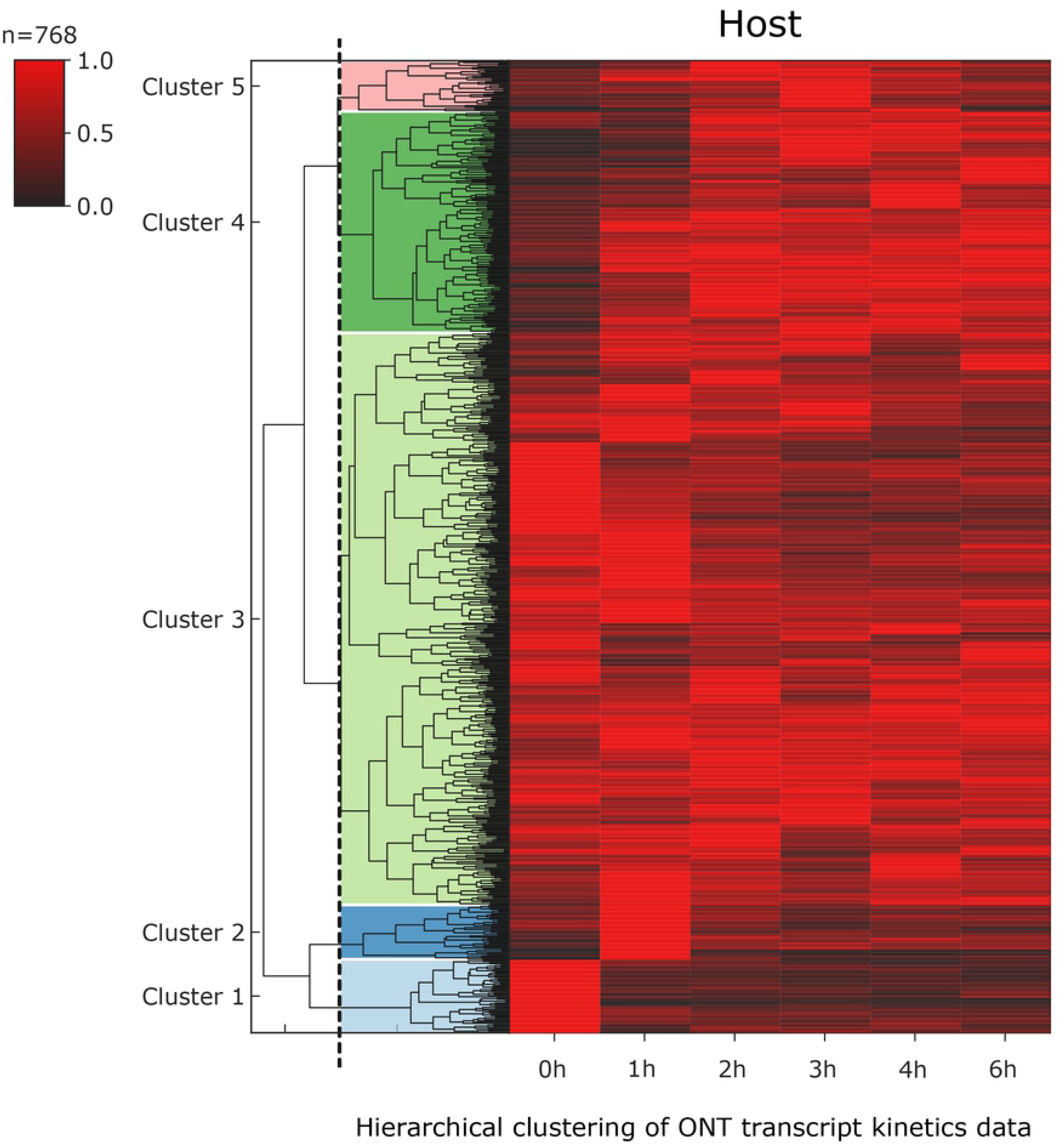
**Heatmap representation of the five distinct host gene clusters.**

In further analyses, we assessed expression patterns of the best characterized gene clusters using gene ontology (GO; **S17 Table**), and found significant overrepresentation of the GO process “regulation of signaling receptor activity” by genes that were highly expressed in the early stages but were not or only slightly expressed in the late phases of infection. Most of the present clusters were not significantly enriched in genes of specific biological processes, although many genes that were upregulated during viral infection were found to play roles in cell division or in “positive regulation of viral life cycle.” Moreover, some genes that were downregulated upon viral infection were annotated to “cell growth” and “mesenchymal differentiation” categories.

## Discussion

SRS has become a standard technique for characterization of transcriptomes^66–67^. Yet comprehensive annotation of transcripts from SRS data remains challenging^68^, and auxiliary methods, such as Northern-blotting and rapid amplification of cDNA ends analyses are too laborious for investigations of large transcriptomes. LRS allows determination of full-length transcripts in single reads without computational inference, and thus distinguishes between transcript isoforms, polycistronic RNAs, and transcript overlaps with ease^69^. In studies at the Moss laboratory, viral transcriptomes were analyzed using Illumina sequencing, and large numbers of TSSs, TESs, and TISs were identified^13, 16, 20^. These studies did not, however, ascertain combinations of TSSs and TESs in transcripts. In this study, we employed two LRS platforms (PacBio and ONT) and sequenced cDNA and dRNA using various library preparation methods for surveying the temporal transcriptional activities of VACV genes. We also analyzed the effects of viral infection on host gene expression. We developed pipelines for profiling long-read RNA sequencing data. In total, RNA molecules may be much more numerous than the transcripts annotated by the LoRTIA toolkit, reflecting high heterogeneity of transcript ends, especially in chaotic genomic regions, which hamper efforts to identify transcripts using bioinformatic techniques. The intermediate and late transcripts are particularly polymorphic in length, but variations in TSS and TES positions of early transcripts also increased at late stages of infection. Generally, canonical transcripts are significantly overrepresented among isoforms encoded in regular genomic regions. Apart from canonical transcripts, VACV genes typically encompass shorter genes with N-terminal truncated ORFs, which may encode shorter polypeptides, likely with altered effector functions. Moreover, longer transcript variants often incorporate uORFs, which may regulate the expression of downstream genes^33^. The genomic organization of poxviruses basically differs from that of herpesviruses. First, poxvirus E genes are gathered at the genome termini, whereas I and L genes are located in central parts of the genome. In addition, proportions of convergent and divergent orientations are low compared with tandemly-oriented genes. In addition, unlike herpesviruses, VACV DNA encodes numerous TSS and TES isoforms. Specifically, whereas prototype herpesvirus transcripts form co-terminal TES clusters, adjacent pox genes produce large numbers of alternative TESs. In VACV, heterogeneity of TESs exceeds that of TSSs, whereas the opposite is the case in herpesviruses. The general presence of within-ORF TSSs is also unique to VACV. Accordingly, using ribosome profiling assays, Yang et al detected several downstream methionine residues and mapped these within main ORFs^13^. We detected numerous TSSs upstream of TISs, thereby providing transcriptional evidence of potential truncated proteins. We also detected long 5’-UTR isoforms containing uORFs in most VACV genes, providing a potential source of micro peptides with regulatory functions, as demonstrated previously in Kaposi’s sarcoma virus^70^. The observed transcript diversity of VACV may, however, represent transcriptional noise from cryptic promoters and/or error-prone transcriptional machinery. Moreover, the majority of transcripts contain multiple translationally active ORFs, and most isoforms contain unique combinations of ORFs and uORFs, suggesting that this transcriptional diversity is functional. Hence, further studies are needed to demonstrate the biological significance of these RNA molecules. We provide evidence that poly(A) signals are frequently transmitted by RNA polymerase without downstream cleavage of transcripts. This process results in the production of polycistronic RNAs from tandem genes and asRNAs from convergent gene pairs. The ensuing stochastic transcription initiation and termination in poxviruses represents a unique gene expression system, which has not yet been described in other well-characterized organisms. No previous studies show translation of polycistronic RNA molecules from downstream genes in poxviruses. Therefore, we can raise the question concerning the function of these multigenic transcripts. To explain this phenomenon and read-through transcription in herpesviruses^71–72^, we hypothesize that transcriptional overlaps are sites of transcriptional interference. Similarly, these interactions have been suggested to represent a genome-wide regulatory system, and the term transcriptional interference networks^73^ (TINs) was coined previously. We report sources of potential error of RT, PCR, and bioinformatic techniques, and propose strategies for addressing these. In particular, we applied very strict criteria for accepting sequencing reads as transcripts so that a large fraction of excluded reads could represent existing transcripts.

Finally, our analyses of uninfected and infected CV-1 cells revealed 758 transcript isoforms, of which only 15% match transcript annotations of the closest relative *C. sabaeus*. In line with previous studies^15, 74^, VACV transcripts were immediately increased at 1h p.i., followed by a rapid decrease in host RNA quantities, reflecting degradation of cellular transcripts by viral decapping enzymes^75–76^. Although partially degraded RNAs are captured by long-read sequencing^77^, we observed relatively constant mean mapped read lengths from host DNA before and during viral infection. The lack of mean read-length shortening could follow low expression of late decapping enzymes (D10) or rapid turnover of degraded RNAs.

## Methods

### Cells and viruses

CV-1 African green monkey kidney fibroblast cells were obtained from the American Type Culture Collection. Cells were plated at a density of 2 × 10^6^ cells per 150-cm^2^ tissue culture flask and were cultured in RPMI 1640 medium (Sigma-Aldrich, St. Louis, MO) supplemented with 10% fetal bovine serum and antibiotic-antimycotic solution (Sigma-Aldrich). CV-1 cells were incubated at 37°C in a humidified atmosphere containing 5% CO_2_ until confluent. Subsequently, about 2.6 × 10^7^ cells were rinsed with serum free RPMI medium immediately prior to infection with the vaccinia virus (VV) WR strain diluted in serum free RPMI medium.

### Infection

Cells were infected with 3-ml aliquots of VV at a multiplicity of infection of 10/cell, and were incubated at 37°C in an atmosphere containing 5% CO_2_ for 1 h with brief agitation at 10 min intervals to redistribute virions. Three-mL aliquots of complete growth medium (RPMI + 10% FBS) were then added to tissue culture flasks and the cells were incubated at 37°C for 1, 2, 3, 4, 6, 8, 12, and 16 h in a humidified atmosphere containing 5% CO_2_. Experiments were performed in triplicate. Following incubation, media were removed and cells were rinsed with serum free RPMI 1640 medium and were subjected to three cycles of freeze–thawing. Cells were finally scraped into 2-ml aliquots of PBS and were stored at −80°C until use.

### RNA purification

Total RNA was extracted from infected cells at various stages of viral infection from 1–12h p.i. using Macherey-Nagel RNA kits according to the manufacturer’s instructions. Polyadenylated RNA fractions were isolated from total RNA samples using Oligotex mRNA Mini Kits (Qiagen) following the Oligotex mRNA Spin-Column Protocol. We used Ribo-Zero Magnetic Kit H/M/R (Illumina) to remove rRNAs from total RNA samples for analyses of non-polyadenylated RNAs.

### PacBio RSII & Sequel sequencing

#### cDNA synthesis

Copy DNAs were generated from polyA(+) RNA fractions using SMARTer PCR cDNA Synthesis Kits (Clontech) according to PacBio Isoform Sequencing (Iso-Seq) using Clontech SMARTer PCR cDNA Synthesis Kit and No Size Selection protocols. Samples from different infection time points (1-, 4-, 8-, and 12-h p.i.) were mixed for RSII library preparation, although samples from 1-, 2-, 3-, 4-, 6-, and 8-h p.i. were used individually to produce libraries for Sequel sequencing. Samples of cDNA were prepared from rRNA-depleted RNA mixtures from 1-, 4-, 8-, and 12-h time points using modified random hexamer primers (**S18 Table**) instead of using the oligo(d)T-containing oligo reagent provided in the SMARTer Kit. Samples were used for SMRTbell template preparation using the PacBio DNA Template Prep Kit 1.0.

#### Purification of samples

Bead-based cDNA purification (Agencourt AMPure XP; Beckman Coulter) was applied after each enzymatic step.

#### SMRTbell library preparation and sequencing

The detailed version of the template preparation protocol is described in our earlier publication^78^. Briefly, primer annealing and polymerase binding reactions were performed using the DNA Sequencing Reagent Kit 4.0 v2 with DNA Polymerase P6 for the RSII platform, whereas the Sequel Sequencing Kit 2.1 and Sequel DNA Polymerase 2.0 were applied for sequencing on the Sequel. Polymerase-template complexes were bound to magbeads prior to loading into the PacBio instrument. Reactions were then performed using The PacBio’s MagBead Kit (v2). Finally, 240-or 600-min movies were captured using the RSII and Sequel machines, respectively. One movie was recorded for each SMRTcell.

### ONT MinION platform – cDNA sequencing

#### 1D cDNA library preparation

PolyA(+) RNA fractions were used for cDNA sequencing on the ONT MinION device. RNAs from different infection time points were converted to cDNAs according to the ONT 1D strand-switching cDNA ligation protocol (Version: SSE_9011_v108_revS_18Oct2016). An RNA mixture containing equal amounts of RNA from 1-, 2-, 3-, 4-, 6-, 8-, 12-, and 16-h p.i. were prepared for sequencing, along with separate RNA samples from each time point. Libraries were generated using the above mentioned 1D ligation kit and protocol, the Ligation Sequencing 1D kit (SQK-LSK108, Oxford Nanopore Technologies), and the NEBNext End repair/dA-tailing Module NEB Blunt/TA Ligase Master Mix (New England Biolabs) according to the manufacturer’s instructions. Briefly, polyA(+)-selected RNAs were converted to cDNAs using Poly(T)-containing anchored primers [(VN)T20; Bio Basic, Canada], dNTPs (10 mM, Thermo Scientific), Superscript IV Reverse Transcriptase Kit (Life Technologies), RNase OUT (Life Technologies), and strand-switching oligonucleotides with three O-methyl-guanine RNA bases (PCR_Sw_mod_3G; Bio Basic, Canada). The resulting double-stranded cDNAs were amplified by PCR using KAPA HiFi DNA Polymerase (Kapa Biosystems), Ligation Sequencing Kit Primer Mix (provided by the ONT 1D Kit), and a Veriti Thermal Cycler (Applied Biosystems). The NEBNext End repair/dA-tailing Module (New England Biolabs) was used to blunt and phosphorylate cDNA ends, and the NEB Blunt/TA Ligase Master Mix (New England Biolabs) was used for adapter (supplied in the 1D kit) ligation.

Size selection: PCR products from mixed RNA samples were size-selected manually and were then run on Ultrapure Agarose gels (Thermo Fischer Scientific). Subsequently, fragments of greater than 500-bp were isolated using Zymoclean Large Fragment DNA Recovery Kits (Zymo Research). Barcoding: Individually sequenced cDNAs were barcoded using a combination of the following ONT protocols: the 1D protocol was used until the first end-preparation step, then we switched to the 1D PCR barcoding (96) genomic DNA (SQK-LSK108) protocol (version: PBGE96_9015_v108_revS_18Oct2016, updated 25/10/2017), followed by the barcode ligation step using the ONT PCR Barcoding Kit 96 (EXP-PBC096). Barcode adapters were ligated to end-prepped samples using Blunt/TA Ligase Master Mix (New England Biolabs).

#### Library preparation from Cap-selected samples

For more accurate definitions of TSSs of full-length RNA molecules, we followed the 5’-Cap-specific cDNA generation protocol combined with the ONT 1D cDNA library preparation method. We produced cDNAs from a mixture of total RNA samples (1-, 2-, 3-, 4-, 6-, 8-, 12-, and 16-h p.i.) using TeloPrime Full-Length cDNA Amplification Kits (Lexogen). Amplified PolyA- and Cap-selected samples were then used for library preparation following the ONT 1D strand-switching cDNA ligation method (ONT Ligation Sequencing 1D kit). Samples were then end-repaired (NEBNext End repair/dA-tailing Module) and ligated to ONT 1D adapters (NEB Blunt/TA Ligase Master Mix).

#### Purification of cDNAs

Samples were purified using Agencourt AMPure XP magnetic beads (Beckman Coulter) after enzyme reaction steps during library preparation.

#### Sequencing on the MinION device

ONT cDNA libraries and Cap-selected samples were loaded onto 3 and 2 ONT R9.4 SpotON Flow Cells for sequencing, respectively. Sequencing runs were performed using MinKNOW.

### ONT MinION platform – dRNA sequencing

To avoid probable amplification-based biases, we used the ONT PCR-free direct RNA (dRNA) sequencing (DRS) protocol (Version: DRS_9026_v1_revM_15Dec2016). A mixture containing PolyA(+) RNAs from eight time points (1-, 2-, 3-, 4-, 6-, 8-, 12-, and 16-h p.i.) was used to generate the library. RNA samples, the oligo(dT)-containing adapter (ONT Direct RNA Sequencing Kit; SQK-RNA001), and T4 DNA ligase (2M U/mL; New England BioLabs) were mixed and incubated for 10 min, and first-strand cDNAs were then prepared using SuperScript III Reverse Transcriptase enzyme (Life Technologies) according to the DRS protocol. The RMX sequencing adapter was ligated to the samples with NEBNext Quick Ligation Reaction Buffer and T4 DNA ligase (New England Bio Labs). Samples were cleaned using AMPure XP beads (Beckman Coulter) after enzymatic steps. Beads were handled before use with RNase OUT (40 U/μL; Life Technologies; 2 U enzyme/1 μL bead). The dRNA library was run on a R9.4 SpotON Flow Cell. Runs were performed using MinKNOW.

### Quantitative reverse transcription PCR

Quantitative reverse transcription-PCR (qRT-PCR; Rotor-Gene Q; Qiagen) was performed to check the specificity of cDNA products derived from Cap-Seq library preparation, and to validate the antisense expression within A11R and A18R genomic loci. Briefly, RT was performed with 20-ng total RNA samples, SuperScript IV reverse transcriptase, and oligo(dT) primers. RT products were then amplified using ABsolute qPCR SYBR Green Mix (Thermo Fisher Scientific) with gene specific primers (**S18 Table**)

### Nucleic acid quality and quantity checking

Concentrations of the reverse-transcribed and adapter-ligated RNAs were measured using a Qubit 2.0 Fluorometer (Life Technologies, **S19 Table**). An Agilent Bioanalyzer 2100 was used for quality control of RNA samples.

### Data analysis and visualization

#### Generation of consensus sequences from the PacBio dataset

ROI reads were created from the RSII raw data using the RS_ReadsOfInsert protocol (SMRT Analysis v2.3.0) with the following settings: Minimum Full Passes = 1, Minimum Predicted Accuracy = 90, Minimum Length of Reads of Insert = 1, Maximum Length of Reads of Insert = No Limit. ROIs from the Sequel dataset were generated using SMRT Link5.0.1.9585.

#### ONT dataset–basecalling

MinION base calling was performed using the ONT Albacore software v.2.0.1 (Albacore, RRID:SCR_015897).

#### Mapping and annotation of TSS and TES positions and transcripts

PacBio ROIs and ONT raw reads were aligned to the reference genome of the virus (LT966077.1) and to that of the host cell (*Chlorocebus sabaeus*; GenBank assembly accession: GCA_000 409795.2 [latest]; RefSeq assembly accession: GCF_000 409795.2 [latest]) using minimap2 aligner (version 2.13) with the options -ax splice -Y -C5–cs. After applying the LoRTIA toolkit (https://github.com/zsolt-balazs/LoRTIA), the determined 5’ and 3’ ends of transcripts and detected full-length reads were mapped. For analyses of transcript dynamics, we included only transcripts that were validated by two different library preparation techniques (PacBio Iso-Seq & MinION 1D or PacBio Iso-Seq and Lexogen TeloPrime or MinION 1D and Lexogen TeloPrime). Output bed files from cDNA sequencing were visualized using Circos plot79 (**Fig 2**).

### Transcript nomenclature

Novel transcripts and ORFs were named using a scheme that incorporates existing ORF names, according to the common (Copenhagen) HindIII fragment letter/number-based nomenclature46. Previously described ORFs were not renamed and our scheme allowed for future additions of novel transcripts. Multicistronic transcripts were named for all contributed ORFs, and the most upstream ORF was listed first. Non-coding transcripts are identified with the prefix “nc,” and complex transcripts carry the prefix “c.” When a non-coding transcript is antisense to the ORF for which it was named, an “as” prefix was added. Names of length isoforms, e.g., TSS and TES isoforms, end with “l” for long, “s” for short, or “AT” for alternative termination (**S1 Note**). Isoform categories are presented in **S12 Figure.**

### Calculation of relative expression levels of viral transcripts

Criteria were set to count relative viral expression values. Specifically, only transcript isoforms annotated by LoRTIA were used for the calculation. In further analyses, criteria were extended to include only transcripts that were represented by at least five independent full-length reads.

Normalized relative expression ratios at a given time point of infection (R_t_) were calculated using the following formula: 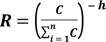, where n = number of different transcript isoforms in a given sample, C = the read count of a given transcript at a given time point, and 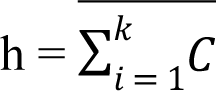 returns the mean C value of a given transcript at time points k = 1, 2, 4, 6, or 8 h. R_t_-values were clustered hierarchically using the Instant Clue statistic program^80^ to determine numbers of transcript isoform clusters. The Euclidian metric was used to cluster rows in the data frame and thereafter k-means hierarchical clustering was conducted with 100000 repetitions. Means of each cluster were presented in BioVinci 1.1.5. Transcripts with at least five full-length reads were considered in cluster analyses. For functional analyses of transcript isoform coding of known VACV proteins, we used descriptions from the UniProt database (https://www.uniprot.org/).

### Analysis of the expression dynamics of VACV ORFs

Normalized relative expression ratios (R_t_ -values) of ORFs at given time points of infection were calculated as described in the previous section with minor modifications. Briefly, R_t_ values of the ORFs were calculated by summing R_t_ values of transcript isoforms that harbor the same ORFs, with the exception of polycistronic transcripts carrying the given ORF in a downstream position. These transcripts are not translated.

### Prediction of *cis*-regulatory sequences

Sequences at 100-nt upstream of a given TSS were extracted and an in-house script based on the GPMiner promoter prediction tool^81–82^ was used for analyses of upstream cis-regulatory sequences of novel transcripts. The general settings of the algorithm were as follows: the eukaryotic promoter database of GPMiner was used to search exact matches without gaps or substitutions. The Vaccinia early promoter motif^15–16, 20^ and late promoter motif^42^ were picked and visualized using the Geneious R10 Motif Finder tool, allowing one substitution in the string. Promoter statistics were calculated using Mann-Whitney U tests with two-tailed p-values.

### Structural analysis of the host transcriptome

Because the host genome assembly was unavailable at the time of this work, reads were aligned to the viral genome and to the genome of *Chlorocebus sabaeus* (NCBI Assembly accession: GCF_000409795.2), which is a close relative of the host *Chlorocebus aethiops*. For analyses of the host transcriptome, data from uninfected and 1-, 2-, 3-, 4-, 6-, 8-, and 12-h p.i. datasets from the MinION sequencing and 1-, 2-, 3-, 4-, 6-, and 8-h p.i. datasets from the Sequel sequencing were used. The remaining samples were pooled with mammalian RNAs from other species. To categorize transcript isoforms of the CV-1 cell line, we used the ti.py script with the transcript annotation files generated by LoRTIA (https://github.com/zsolt-balazs/LoRTIA). Isoforms were determined according to the positions of their TSSs, TESs, splice junctions, and orientations relative to the previous annotation (NCBI Assembly accession: GCF_000409795.2). Isoform categories are presented in **S12 Figure.**

### Time-course analysis of gene expression in the host *C. aethiops* cell line

Raw fastq files of MinION long reads were aligned to a joint host genome because there is no reference genome for *C. aethiops*. The reference genome of *C. sabeus* GCF_000409795.2 was used. We excluded MAPQ = 0 and secondary and supplementary alignments from downstream analyses. Primary alignments mapping to the pox genome were counted separately. Reads that were aligned to the host genome were associated with host genes according to the *C. sabeus*_1.1_top_level.gff3 genome coordinates. Only reads that matched the exon structure of reference genes (using a ±5-bp window for matching exon start and end positions) were counted. Gene counts were normalized to total valid read counts that were mapped to the host genome to identify abundant housekeeping genes that were not influenced by virus–host interactions during the experiment. Geometric means were calculated for the top 16 housekeeping genes (*RPS18*, *RPL27*, *EEF1A1*, *CST3*, *MYL12B*, *RPL23*, *RPS5*, *RPL17*, *UBL5*, *RPS7*, *TPM1*, *RPL5*, *ATP5A1*, *LOC103217585*, *RPL27A*, and *RPS14*) and a coefficient of variation (CV) of <0.2 was used to normalize gene counts. Our criteria for statistical analyses of gene expression included >60 normalized total gene (highly expressed) counts for the six-time-point data. To classify genes by their gene expression profiles in the examined time series, we transformed normalized gene counts to a relative scale where the highest expression time point had a value of 1.0 (100%). We performed clustering using the k-means algorithm of the basic statistics package of R (version 3.5). The optimal number of clusters was determined using Calinski-Harabasz criteria^83^ with the cascade KM algorithm of the vegan (version 2.5-4) R package. Clusters were visualized using the heatmap (version 1.0.2) R package. Because few host gene reads failed to enable characterization of host gene expression at the later time points (8- and 12-h p.i.), host gene expression was only analyzed until 6 h after infection. Based on the Calinski-Harabasz criteria, the optimal number of clusters was five for our dataset (**S13 Figure**). Accordingly, we clustered the genes into five subcategories according to normalized gene expression profiles. However, because clustering forces genes into categories, we define the subsets of genes with the most typical expression profiles for their category. Thus, in each cluster we plotted mean expression levels for different sampling times using ggplot2 (version 3.1.0, stat_smooth algorithm with the loess method). According to these plots, we identified four categories for which gene expression profiles changed during the experiment (**Fig 14**).

Based on the expression profiles of these different categories, we calculated scores for each gene in every cluster by subtracting mean relative gene expression levels of low expression sampling points from the means of high expression points. We identified the most typical characteristic genes in all clusters falling within the range between the top score and the top score–1 SD (**S20 Table**). Using the identified subset of genes, we performed overrepresentation analyses for the most characteristic genes in each cluster using 768 highly expressed genes as references with the PANTHER (version 14.1 using the 2018_04 dataset release)^84^ software tool. We then analyzed GO biological processes with false discovery rates (FDR) of less than 1.

## Data availability

Relevant data are within the manuscript and its supporting information files. In addition, long-read sequencing datasets are available in European Nucleotide Archive (ENA) under the accession number PRJEB26434 and PRJEB26430. The complete map of the VACV transcriptome is available at FigShare: 10.6084/m9.figshare.10191230 as a Geneious file.

## Acknowledgements

This study was supported by National Research, Development and Innovation Office (OTKA) grants K 128247 to ZBo and FK 128252 to DT. DT was also supported by the Eötvös Scholarship of the Hungarian State. IP was supported by the UNKP-19-3-SZTE-243 New National Excellence Program of the Ministry for Innovation and Technology. The project was also supported by the NIH Centers of Excellence in Genomic Science Center for Personal Dynamic Regulomes [5P50HG00773502] to MS. The funders had no role in study design, data collection and analysis, decision to publish, or preparation of the manuscript. The authors thank Annamária Gáspárné (Veterinary Diagnostic Directorate of the NFCSO) for laboratory assistance.

## Author Contributions

D.T., M.S., and Z.Bo. designed the study. Z.M., T.K. and N.M. provided additional input into study design. D.T. carried out the PacBio sequencing. D.T. and Z.C. performed the MinION sequencing. D.T., I.P., B.D., and Z.C. performed all other experiments. D.T., I.P., Z.Ba., N.M., Z.M., T.K. and Z. Bo. analyzed the data. D.T., I.P., N.M. and Z.Bo drafted the manuscript. D.T. N.M and I.P. made the figures. D.T. and Z.Bo. wrote the final version of the manuscript. All authors read and approved the final paper.

## Abbreviations

asRNA: antisense RNA
cxRNA: complex RNA
dRNA: direct RNA (sequencing)
E: early (gene/transcript)
I: intermediate (gene/transcript)
Iso-Seq: isoform sequencing
IE: immediate-early (gene/transcript)
L: late gene/transcript (gene/transcript)
lncRNA: long non-coding RNA
LRS: long-read sequencing
ncRNA: non-coding RNA
ONT: Oxford Nanopore technologies
ORF: open reading frame
Ori: replication origin
PAT: polyadenylation tail
PacBio: Pacific Bioscience
ROI: Reads of Insert
SMRT: single molecule, real-time
sncRNA: short non-coding RNA
SRS: short-read sequencing
TES: transcription end sites [equivalent of poly(A) sites (PAS)]
TIS: translation initiation site
TSS: transcription start site
UTR: untranslated region
VACV: vaccinia virus

---

**S1 Figure. The B12R-B13R-B15R region of the VACV genome as a typical example of a regular transcript region in the VACV transcriptome; a.** illustration of polycistronic transcripts within the B12R-B15R region; **b.** examples of non-coding transcripts; **c.** representation of monocistronic transcripts and 5’- and 3’-isoforms.

**S2 Figure. 5’-untranslated region (UTR) length distributions of transcript variants;** the column diagram and the violin plot show 5’-UTR lengths of VACV transcripts ranging from 2 to 400 nts.

**S3 Figure. Fully overlapping complex transcripts. A.** A8R-A9L genomic loci. **B**. E10R-E11L region. **Color legend:** yellow: annotated ORFs; blue: transcripts, determined by this study; magenta/purple: cxRNAs

**S4 Figure. Shared transcription start and end site positions across the VACV genome. A.** TSS sites shared by ≥ 1 transcript isoforms; the X axis represents genome positions and the Y axis shows the number of transcript isoforms. **B.** TES positions shared by ≥ 1 RNA isoforms; the X axis represents the genome positions and the Y axis shows the number of transcript isoforms.

**S5 Figure. Transcript isoforms;** the X-axis represents genomic coordinates of the first ATGs of the ORFs. The number of LoRTIA-annotated transcript isoforms of given genes are shown on Y-axes.

**S6 Figure. Genome-wide kinetics of VACV transcription start and end sites.** TSSs (panel A) and TESs (panel **B**) were determined using the LoRTIA software suite in each sample. Blue dashes represent TSSs/TESs on the forward strand, while red dashes represent TSSs on the opposite DNA strand. Orange rectangles represent the ORFs.

**S7 Figure. Gene expression curves;** expression patterns of the five distinct viral gene clusters are illustrated in the line graph.

**S8 Figure. Heatmap representation of the VACV ORFs;** Rt values of given ORFs were derived from the sum of viral transcript isoforms of that ORF. Rows represent changes in relative RNA expression. Red rectangles indicate high relative expression values and black rectangles indicate low relative expression values.

**S9 Figure. The length of 3’-UTRs of the alternatively terminated transcript isoforms.**

**S10 Figure. Mapped length distributions of MinION and Sequel sequencing reads;** the upper halves of the density plots of VACV reads from MinION sequencing are not shown. MinION is well-suited for sequencing of smaller VACV RNAs, which are lost during library preparation for the Sequel platform.

**S11 Figure. Transcript and UTR lengths of the host** (*C. aethiops*). **A**. Length of transcripts in the uninfected and each p.i. samples; **B**-**C**.; length of the 5’ and 3’ UTRs were calculated using ti.py in uninfected and each p.i. samples. Letter ‘M’ following the sample name indicates MinION sequencing, and the letter ‘S’ indicates Sequel sequencing. Horizontal lines in the box plots represent median transcript length of the given samples. Transcripts were annotated using the LoRTIA software suit.

**S12 Figure. The transcript isoform categories used in this study and their abbreviations.**

**S13 Figure. Optimal number of clusters based on Calisnki-Harabasz (CH) criterion.** The plot on the left shows how each of the genes are partitioned with an increasing number of clusters. On the right, the maximum CH index (for 5 clusters) is shown.

**S1 Table. Statistics**

**S2 Table. Summary table of average and median values of coding sequences of various viruses.** Abbreviations: VACV: Vaccinia virus; HSV: Herpes-simplex virus type-1; PRV: Pseudorabies virus; CMV: Human Cytomegalovirus; EBV: Epstein-Barr virus; ASFV: African swine fever virus; AcMNPV: *Autographa californica* multicapsid nucleopolyhedrovirus

**S3 Table. Summary table of the transcripts identified by this study. A.** Genomic position of the 8,191 transcripts identified by the LoRTIA toolkit. **B.** Names and position of the 1,480 VACV transcripts identified by the LoRTIA tool under strict criteria. **C.** List of additional 308 transcripts (from the 8,191 list) that were identified by the LoRTIA pipeline; these contain ORF but do not meet the strict criteria. **D.** 23 transcripts that do not meet the LoRTIA criteria.

**S4 Table. List of the novel intergenic genes;** previously published corresponding TSS positions, ORF information, and the distance of the TSSs of novel genes from the predicted promoters are also summarized.

**S5 Table. Comparison of the TSS positions of newly identified embedded genes and previously published translation initiation site (TIS) coordinates**

**S6 Table. List of the VACV genes with longer TSS isoforms than the canonical transcripts;** length of ORFs within the longer UTR variants are presented with differences between main and longer isoforms and numbers of ORFs within longer variants.

**S7 Table. List of the monocistronic VACV transcripts.** Black letters: LoRTIA transcripts with strict criteria; Blue letters: LoRTIA transcripts; Red letters: additional monocistronic transcripts.

**S8 Table. List of the complex RNAs.**

**S9 Table. Comparison of TSS positions from our experiments and those from previous studies. A**. Summary table of the previously publishes TSSs and the TSSs of this study. **B.** References of the TSSs described in earlier studies.

**S10 Table. List of genes with newly identified TATA boxes; t**he table contains the genomic location of genes, TATA-box sequences and their distances from the TSSs.

**S11 Table. List of the convergent, divergent and parallel gene-pairs of VACV** Types of transcriptional overlaps are presented as follows: no overlap, ORF-ORF overlap, transcript-ORF overlap, transcript-transcript overlap.

**S12 Table. List of the polycistronic transcripts;** transcripts were identified using the LoRTIA toolkit under strict criteria.

**S13 Table. VACV gene expression; A.** Relative expression values of the viral genes; the genes are arranged according to their kinetic clusters. **B**. Detailed information about the function of examined VACV genes.

**S14 Table. Transcriptional start and end sites of the host (*C. aethiops*) RNAs;** read counts, distances between GC-, CAAT- and TATA-boxes and TSSs, the sequence of these features, the distance between polyadenylation signals and TESs, and sequences of ±50-nt regions of TESs are shown. The letter ‘M’ following the sample name indicates MinION sequencing and the letter ‘S’ refers to Sequel sequencing. The TSS and TES sequences were identified using the LoRTIA software suit.

**S15 Table. Introns of the host transcripts**; splice donor and acceptor positions are shown with read counts and the sequences of the splice junctions. Letter ‘M’ following the sample name indicates MinION sequencing, while letter ‘S’ refers to Sequel sequencing. The introns were determined using the LoRTIA pipeline.

**S16 Table. Transcript isoforms of host cells;** read counts, category abbreviations, length of transcripts, and length of 5’ and 3’ UTRs are shown. Abbreviation of categories is defined in **S12 Figure.** Transcript isoforms were determined using the LoRTIA toolkit and were categorized using the ti.py script.

**S17 Table. Overrepresentation analysis of host gene expression levels;** the first column contains clusters of host genes and numbers of genes that were characteristic of the cluster (in parentheses). GO biological processes (bold) that had the lowest false discovery rates (FDR) values are presented with numbers of genes in clusters and numbers of genes in the reference dataset (in parentheses). The genes belonging to GO processes are also listed.

**S18 Table. List of primers used in this study.**

**S19 Table. List of the Qubit Assays used for nucleic acid quantitation.**

**S20 Table. Clustering of the 768 most active host genes based on their expression during VACV infection**; relative expression values at each p.i. time point are normalized to the highest value for a given gene. Genes that were most characteristic of each cluster are highlighted in gray.

**S1 Note. Terminology of the listed transcripts.**

**S2 Note**. **Information for downloading and using the Geneious file S3 Note**. **List of transcripts with unknown or non-validated TSSs**.

## References

1. Moss B. Poxviridae. In Knipe DM, Howley PM, Cohen JI, Griffin DE, Lamb RA, Martin MA, Rancaniello VR, Roizman B, editors. Fields virology. Philadelphia: Lippincott Williams & Wilkins; 2013. pp. 2129–2159.

2. Esposito JJ, Sammons SA, Frace AM, Osborne JD, Olsen-Rasmussen M, Zhang M, et al. Genome sequence diversity and clues to the evolution of variola (smallpox) virus. Science. 2006;313:807– 812. doi:10.1126/science.1125134

3. Jacobs BL, Langland JO, Kibler KV, Denzler KL, White SD, Holechek SA, et al. Vaccinia Virus Vaccines: Past, Present and Future. Antiviral Res. 2009;84:1–13. doi: 10.1016/j.antiviral.2009.06.006

4. Garcel A, Crance J, Drillien R, Garin D, Favier A. Genomic sequence of a clonal isolate of the vaccinia virus Lister strain employed for smallpox vaccination in France and its comparison to other orthopoxviruses. J Gen Virol. 2007;88:1906–1916. doi:10.1099/vir.0.82708-0

5. . Moss B. Poxviridae: The viruses and their replication. In Knipe DM, Howley PM, editors. Fields Virology. Philadelphia: Lippincott Williams & Wilkins; 2007. pp. 2905–2946.

6. Yang Z, Moss B. Decoding poxvirus genome. Oncotarget. 2015;30:28513–28514.

7. Wei CM, Moss B. Methylated nucleotides block 5-terminus of vaccinia virus mRNA. Proc Natl Acad Sci USA. 1975;72:318–322.

8. Kates J, Beeson J. Ribonucleic acid synthesis in vaccinia virus. I. The mechanism of synthesis and release of RNA in vaccinia cores. J Mol Biol. 1970;50:1–18.

9. Davison AJ, Moss B. Structure of vaccinia virus early promoters. J Mol Biol. 1989;210:749–769.

10. Davison AJ, Moss B. Structure of vaccinia virus late promoters. J Mol Biol. 1989;210:771–784.

11. Baldick CJ Jr, Keck JG, Moss B. Mutational analysis of the core, spacer, and initiator regions of vaccinia virus intermediate-class promoters. J Virol. 1992;66:4710–4719.

12. Broyles S: Vaccinia Virus Transcription. J Gen Virol. 2003;84:2293–2303.

13. Yang Z, Cao S, Martens CA, Porcella SF, Xie Z, Ma M, et al. Deciphering poxvirus gene expression by RNA sequencing and ribosome profiling. J Virol. 2015 Jul;89(13):6874–86. doi: 10.1128/JVI.00528-15.

14. Assarsson E, Greenbaum JA, Sundström M, Schaffer L, Hammond JA, Pasquetto V, et al. Kinetic analysis of a complete poxvirus transcriptome reveals an immediate-early class of genes. Proc Natl Acad Sci USA. 2008;105(6):2140–5. doi: 10.1073/pnas.0711573105.

15. Yang Z, Bruno DP, Martens CA, Porcella SF, Moss B. Simultaneous high-resolution analysis of vaccinia virus and host cell transcriptomes by deep RNA sequencing. Proc Natl Acad Sci USA. 2010;107(25):11513–8. doi: 10.1073/pnas.1006594107.

16. Yang Z, Reynolds SE, Martens CA, Bruno DP, Porcella SF, Moss B. Expression Profiling of the Intermediate and Late Stages of Poxvirus Replication J Virol. 2011;85(19):9899–9908. doi:10.1128/JVI.05446-11 2011.

17. Honess RW, Roizman B. Regulation of herpesvirus macromolecular synthesis: Sequential transition of polypeptide synthesis requires functional viral polypeptides. Proc Natl Acad Sci USA. 1975;72:1276–1280.

18. Ross L, Guarino LA. Cycloheximide inhibition of delayed early gene expression in baculovirus-infected cells. Virology. 1997;232:105–113.

19. Cooper JA, Moss B. In vitro translation of immediate early, early, and late classes of RNA from vaccinia virus-infected cells. Virology. 1979;96:368–380.

20. Yang Z, Bruno DP, Martens CA, Porcella SF, Moss B. Genome-wide analysis of the 5’ and 3’ ends of vaccinia virus early mRNAs delineates regulatory sequences of annotated and anomalous transcripts. J Virol. 2011;85(12):5897–909. doi: 10.1128/JVI.00428-11.

21. Yuen L, Moss B. Oligonucleotide sequence signaling transcriptional termination of vaccinia virus early genes. Proc Natl Acad Sci USA. 1987;84:6417–6421.

22. Shuman S, Moss B. Factor-dependent transcription termination by vaccinia virus RNA polymerase. Evidence that the cis-acting termination signal is in nascent RNA. J Biol Chem. 1988;263:6220–6225.

23. Ahn BY, Jones EV, and Moss B. Identification of the vacciniavirus gene encoding an 18-kilodalton subunit of RNA polymerase and demonstration of a 5’ poly(A) leader on its early transcript. J Virol. 1990;64:3019–3024.

24. Rubins KH, Hensley LE, Bell GW, Wang C, Lefkowitz EJ, Brown PO, et al. Comparative analysis of viral gene expression programs during poxvirus infection: a transcriptional map of the vaccinia and monkey pox genomes. PLoS One. 2008;3:e2628. doi.org/10.1371/journal.pone.0002628

25. Stern-Ginossar N, Weisburd B, Michalski A, Le VT, Hein MY, Huang SX, et al. Decoding human cytomegalovirus. Science. 2012;338:1088–1093. doi.org/10.1126/science.1227919.

26. Guerra S, López-Fernández LA, Pascual-Montano A, Muñoz M, Harshman K, Esteban M. Cellular gene expression survey of vaccinia virus infection of human HeLa cells. J Virol. 2003;77:6493–6506.

27. Brum LM, Lopez MC, Varela JC, Baker HV, Moyer RW. Microarray analysis of A549 cells infected with rabbitpox virus (RPV): A comparison of wild-type RPV and RPV deleted for the host range gene, SPI-1. Virology. 2003;315:322–334.

28. Tombácz D, Csabai Z, Oláh P, Balázs Z, Likó I, Zsigmond L, Sharon D, et al. Full-Length Isoform Sequencing Reveals Novel Transcripts and Substantial Transcriptional Overlaps in a Herpesvirus. PLoS One. 2016;11(9):e0162868. doi: 10.1371/journal.pone.0162868

29. Moldován N, Tombácz D, Szűcs A, Csabai Z, Snyder M, Boldogkői Z. Multi-Platform Sequencing Approach Reveals a Novel Transcriptome Profile in Pseudorabies Virus. Front Microbiol. 2018; 8:2708. doi.org/10.3389/fmicb.2017.02708

30. Tombácz D, Csabai Z, Szűcs A, Balázs Z, Moldován N, Sharon D, et al. Long-read isoform sequencing reveals a hidden complexity of the transcriptional landscape of Herpes simplex virus type 1. Front Microbiol. 2017;8:1079. doi: 10.3389/fmicb.2017.01079.

31. Tombácz D, Moldován N, Balázs Z, Gulyás G, Csabai Z, Boldogkői M, et al. Multiple Long-Read Sequencing Survey of Herpes Simplex Virus Dynamic Transcriptome. Front Genet. 2019;10:834. doi: 10.3389/fgene.2019.00834.

32. Prazsák I, Moldován N, Balázs Z, Tombácz D, Megyeri K, Szűcs A, et al. Long-read sequencing uncovers a complex transcriptome topology in varicella zoster virus. BMC Genomics. 2018;19(1):873. doi: 10.1186/s12864-018-5267-8.

33. Balázs Z, Tombácz D, Szűcs A, Csabai Z, Megyeri K, Petrov AN, et al. Long-Read Sequencing of Human Cytomegalovirus Transcriptome Reveals RNA Isoforms Carrying Distinct Coding Potentials. Sci Rep. 2017;7(1):15989. doi: 10.1038/s41598-017-16262-z.

34. Balázs Z, Tombácz D, Szűcs A, Snyder M, Boldogkői Z. Long-read sequencing of the human cytomegalovirus transcriptome with the Pacific Biosciences RSII platform. Sci Data. 2017;4:170194. doi: 10.1038/sdata.2017.194.

35. Tombácz D, Balázs Z, Csabai Z, Moldován N, Szűcs A, Sharon D, et al. Characterization of the Dynamic Transcriptome of a Herpesvirus with Long-read Single Molecule Real-Time Sequencing. Sci Rep. 2017;7:43751. doi: 10.1038/srep43751.

36. Boldogkői Z, Moldován N, Balázs Z, Snyder M, Tombácz D. Long-Read Sequencing - A Powerful Tool in Viral Transcriptome Research. Trends Microbiol. 2019;27(7):578–592. doi: 10.1016/j.tim.2019.01.010.

37. Balázs Z, Tombácz D, Csabai Z, Moldován N, Snyder M, Boldogkői Z. Template-switching artifacts resemble alternative polyadenylation. BMC Genomics. 2019;20(1):824. doi: 10.1186/s12864-019-6199-7.

38. Zhu YY, Machleder EM, Chenchik A, Li R, Siebert PD. Reverse transcriptase template switching: a SMART approach for full-length cDNA library construction. Biotechniques. 2001;30: 892–7. doi.org/10.2144/01304pf02

39. Miyamoto M, Motooka D, Gotoh K, Imai T, Yoshitake K, Goto N, et al. Performance comparison of second-and third-generation sequencers using a bacterial genome with two chromosomes. BMC Genomics. 2014;15:699. doi: 10.1186/1471-2164-15-699

40. Tombácz D, Csabai Z, Oláh P, Havelda Z, Sharon D, Snyder M, et al. Characterization of novel transcripts in pseudorabies virus. Viruses. 2015;7:2727–44. doi:10.3390/v7052727

41. Yang Z, Martens CA, Bruno DP, Porcella SF, Moss B. Pervasive initiation and 3’-end formation of poxvirus postreplicative RNAs. J Biol Chem. 2012;287(37):31050–60. doi: 10.1074/jbc.M112.390054.

42. Engelstad M, Howard ST, Smith GL. A constitutively expressed vaccinia gene encodes a 42-kDa glycoprotein related to complement control factors that forms part of the extracellular virus envelope. Virology. 1992;188(2):801–10.

43. Lee-Chen GJ, Bourgeois N, Davidson K, Condit RC, Niles EG. Structure of the transcription initiation and termination sequences of seven early genes in the vaccinia virus HindIII D fragment. Virology. 1988;163(1):64–79.

44. Schmitt JF, Stunnenberg HG. Sequence and transcriptional analysis of the vaccinia virus HindIII I fragment. J Virol. 1988;62(6):1889–97.

45. Tengelsen LA, Slabaugh MB, Bibler JK, Hruby DE. Nucleotide sequence and molecular genetic analysis of the large subunit of ribonucleotide reductase encoded by vaccinia virus. Virology. 1988;164(1):121–31.

46. Rosel JL, Earl PL, Weir JP, Moss B. Conserved TAAATG sequence at the transcriptional and translational initiation sites of vaccinia virus late genes deduced by structural and functional analysis of the HindIII H genome fragment. J Virol. 1986;60(2):436–49.

47. Broyles SS, Moss B. Homology between RNA polymerases of poxviruses, prokaryotes, and eukaryotes: nucleotide sequence and transcriptional analysis of vaccinia virus genes encoding 147-kDa and 22-kDa subunits. Proc Natl Acad Sci USA. 1986;83(10):3141–5.

48. Larkin J, Henley RY, Jadhav V, Korlach J, Wanunu M. Length-independent DNA packing into nanopore zero-mode waveguides for low-input DNA sequencing. Nat Nanotechnol. 2017;12:1169– 1175. doi:10.1038/nnano.2017.176

49. Prazsák I, Tombácz D, Szűcs A, Dénes B, Snyder M, Boldogkői Z. Full Genome Sequence of the Western Reserve Strain of Vaccinia Virus Determined by Third-Generation Sequencing. Genome Announc. 2018;6(11):e01570–17. doi: 10.1128/genomeA.01570-17

50. Ardui S, Ameur A, Vermeesch JR, Hestand MS. Single molecule real-time (SMRT) sequencing comes of age: applications and utilities for medical diagnostics. Nucleic Acids Res. 2018;46(5):2159–2168. doi: 10.1093/nar/gky066.

51. Moldován N, Tombácz D, Szűcs A, Csabai Z, Balázs Z, Kis E, et al. Third-generation Sequencing Reveals Extensive Polycistronism and Transcriptional Overlapping in a Baculovirus. Sci Rep. 2018;8:8604. doi:10.1038/s41598-018-26955-8

52. Goebel SJ, Johnson GP, Perkus ME, Davis SW, Winslow JP, Paoletti E. The complete DNA sequence of vaccinia virus. Virology. 1990;179(1):247–266. doi:10.1016/0042-6822(90)90294-2

53. Somers J, Pöyry T, Willis AE. A perspective on mammalian upstream open reading frame function. Int J Biochem Cell Biol. 2013;45(8):1690–700. doi: 10.1016/j.biocel.2013.04.020.

54. Calvo SE, Pagliarini DJ, Mootha VK. Upstream open reading frames cause widespread reduction of protein expression and are polymorphic among humans. Proc Natl Acad Sci USA. 2009;106:7507–7512.

55. Matsui M, Yachie N, Okada Y, Saito R, Tomita M. Bioinformatic analysis of post-transcriptional regulation by uORF in human and mouse. FEBS Lett. 2007;581:4184–4188.

56. Scholz A, Eggenhofer F, Gelhausen R, Grüning B, Zarnack K, Brüne B. uORF-Tools-Workflow for the determination of translation-regulatory upstream open reading frames. PLoS One. 2019;14(9):e0222459. doi: 10.1371/journal.pone.0222459.

57. Senkevich TG, Bruno D, Martens C, Porcella SF, Wolf YI, Moss B. Mapping vaccinia virus DNA replication origins at nucleotide level by deep sequencing. Proc Natl Acad Sci USA. 2015;112(35):10908–13. doi: 10.1073/pnas.1514809112.

58. Boldogkői Z, Balázs Z, Moldován N, Prazsák I, Tombácz D. Novel classes of replication-associated transcripts discovered in viruses. RNA Biol. 2019;16(2):166–175. doi: 10.1080/15476286.2018.1564468.

59. Smith GL, Chan YS, Kerr SM. Transcriptional mapping and nucleotide sequence of a vaccinia virus gene encoding a polypeptide with extensive homology to DNA ligases. Nucleic Acids Res. 1989;17(22):9051–9062.

60. Brown CK, Turner PC, Moyer RW. Molecular characterization of the vaccinia virus hemagglutinin gene. J Virol. 1991;65(7):3598–3606.

61. Grossegesse M, Doellinger J, Haldemann B, Schaade L, Nitsche A. A Next-Generation Sequencing Approach Uncovers Viral Transcripts Incorporated in Poxvirus Virions. Viruses. 2017;9(10):296. doi:10.3390/v9100296.

62. Vo Ngoc L, Cassidy CJ, Huang CY, Duttke SHC, Kadonaga JT. The human initiator is a distinct and abundant element that is precisely positioned in focused core promoters. Genes Dev. 2017;31(1):6–11. doi: 10.1101/gad.293837.116.

63. Rubins KH, Hensley LE, Relman DA, Brown PO: Stunned Silence: Gene Expression Programs in Human Cells Infected with Monkeypox or Vaccinia Virus. PLoS One. 2011;6(1): e15615. doi: 10.1371/journal.pone.0015615.

64. Bourquain D, Dabrowski PW, Nitsche A: Comparison of host cell gene expression in cowpox, monkeypox or vaccinia virus-infected cells reveals virus-specific regulation of immune response genes Virol J. 2013;10:61. doi: 10.1186/1743-422X-10-61

65. Soday L, Lu Y, Albarnaz JD, Davies CTR, Antrobus R, Smith GL, et al. Quantitative Temporal Proteomic Analysis of Vaccinia Virus Infection Reveals Regulation of Histone Deacetylases by an Interferon Antagonist. Cell Rep. 2019;27(6):1920–1933.e7. doi: 10.1016/j.celrep.2019.04.042.

66. Mortazavi A, Williams BA, McCue K, Schaeffer L, Wold B. Mapping and quantifying mammalian transcriptomes by RNA-Seq. Nat Methods. 2008;5: 621–8. doi: 10.1038/nmeth.1226

67. Djebali S, Davis CA, Merkel A, Dobin A, Lassmann T, Mortazavi A, et al. Landscape of transcription in human cells. Nature. 2012;489: 101–8. doi: 10.1038/nature11233.

68. Steijger T, Abril JF, Pär G Engström, Felix Kokocinski, The RGASP Consortium, Tim J Hubbard et al. Assessment of transcript reconstruction methods for RNA-seq. Nat Methods. 2013;10:1177– 84. doi:10.1038/nmeth.2714

69. Rhoads A, Au KF. PacBio Sequencing and Its Applications. Genomics Proteomics Bioinformatics. 2015;13:278–89. doi.org/10.1016/j.gpb.2015.08.002

70. Kronstad LM, Brulois KF, Jung JU, Glaunsinger BA. Dual short upstream open reading frames control translation of a herpesviral polycistronic mRNA. PLoS Pathog. 2013;9(1):e1003156. doi: 10.1371/journal.ppat.1003156.

71. Tombácz D, Balázs Z, Csabai Z, Snyder M, Boldogkői Z. Long-Read Sequencing Revealed an Extensive Transcript Complexity in Herpesviruses. Front Genet. 2018;9:259. doi: 10.3389/fgene.2018.00259

72. Boldogkői Z, Tombácz D, Balázs Z. Interactions between the transcription and replication machineries regulate the RNA and DNA synthesis in the herpesviruses. Virus Genes. 2019;55(3):274–279. doi: 10.1007/s11262-019-01643-5

73. Boldogkői Z. Transcriptional interference networks coordinate the expression of functionally related genes clustered in the same genomic loci. Front Genet. 2012;3:122. doi: 10.3389/fgene.2012.00122

74. Boone RF, Moss B. Sequence complexity and relative abundance of vaccinia virus mRNA’s synthesized in vivo and in vitro. J Virol. 1978;26(3): 554–69.

75. Parrish S, Moss B. Characterization of a second vaccinia virus mRNA-decapping enzyme conserved in poxviruses. J Virol. 2007;81(23):12973–8. doi: 10.1128/JVI.01668-07

76. Parrish S, Resch W, Moss B. Vaccinia virus D10 protein has mRNA decapping activity, providing a mechanism for control of host and viral gene expression. Proc Natl Acad Sci USA 2007;104(7):2139–44. doi: 10.1073/pnas.0611685104

77. Ozsolak F, Milos PM. RNA sequencing: advances, challenges and opportunities. Nat Rev Genet. 2011;12(2):87–98. doi: 10.1038/nrg2934.

78. Tombácz D, Sharon D, Szűcs A, Moldován N, Snyder M, Boldogkői Z. Transcriptome-wide survey of pseudorabies virus using next- and third-generation sequencing platforms. Sci Data. 2018;5:180119. doi: 10.1038/sdata.2018.119

79. Krzywinski M, Schein J, Birol I, et al. Circos: an information aesthetic for comparative genomics. Genome Res. 2009;19(9):1639–45. doi: 10.1101/gr.092759.109

80. Nolte H, MacVicar TD, Tellkamp F, Krüger M. Instant Clue: A Software Suite for Interactive Data Visualization and Analysis. Sci Rep. 2018;8(1):12648. doi: 10.1038/s41598-018-31154-6

81. Lee TY, Chang WC, Hsu JB, Chang TH, Shien DM. GPMiner: an integrated system for mining combinatorial cis-regulatory elements in mammalian gene group. BMC Genomics. 2012;13:S3. doi:10.1186/1471-2164-13-S1-S3

82. Narang V, Sung WK, Mittal A. Computational modeling of oligonucleotide positional densities for human promoter prediction. Artif Intell Med. 2005;9–10;35(1-2):107-19. doi: 10.1016/j.artmed.2005.02.005

83. Calinski, T. and J. Harabasz. A dendrite method for cluster analysis. Commun Stat. 1974;3(1):1–27.

84. Mi H, Muruganujan A, Casagrande JT, Thomas PD. Large-scale gene function analysis with the PANTHER classification system. Nat Protoc. 2013;8:1551–66. doi:10.1038/nprot.2013.092.

